# Transcriptomic profiling of the adult reptilian dentition sheds light on the genes regulating indefinite tooth replacement

**DOI:** 10.1101/2022.12.23.521841

**Authors:** Joaquin Ignacio Henriquez, Stephane Flibotte, Katherine Fu, Edward Zirui-Wang Li, Joy M. Richman

## Abstract

The aim of this study is to profile the transcriptome of teeth and the surrounding tissues of an adult lizard dentition (*Eublepharis macularius*) that is actively replacing teeth throughout life. Bulk RNAseq was used to compare teeth that are in function versus unerupted, developing teeth and single cell RNA-seq was carried out on jaw segments containing the dental forming tissues. In bulk RNAseq data, we found that functional teeth expressed genes involved in bone and tooth resorption. Indeed, multinucleated odontoclasts were abundant in tissue sections of functional teeth undergoing resorption. Unexpectedly, chemotaxis gene *SEMA3A* was expressed within odontoblasts and in adjacent mesenchyme, confirmed using RNAscope. Semaphorins may be involved in regulating odontoclasts during tooth resorption. The scRNA-seq experiment successfully isolated dental mesenchyme and epithelial cells. We confirmed that some of these genes are expressed in the earliest tooth buds within the tooth forming field. In addition, we found evidence of convergent evolution in the tooth eruption trait. Geckos evolved a means for second generation teeth to communicate with the functional teeth. Instead of a dental follicle inducing an eruption pathway as in the mammal, the gecko and other squamate reptiles use the enamel organ of the successional teeth to trigger tooth resorption of the functional teeth, thus creating an eruption pathway. New molecules such as SEMA3A and SFRP2 may also participate in this process. Future studies on the gecko will uncover the molecular basis of convergent evolution in the dentition.

## Introduction

Tooth replacement is the process by which a functional tooth present in the oral cavity is shed and a successor tooth erupts and takes its place. There is a great diversity across species about the number of times that tooth replacement occurs during the life span of an animal ranging from life-long replacement (Polyphyodonty) to no replacement (Monophyodonty). Most non-mammalian vertebrates including fish, amphibians and reptiles are polyphyodont (Tucker and Fraser 2014). In the mammalian lineage the common pattern is for partial diyphyodonty. In humans there are 20 primary teeth that are replaced with the incisors, canines and premolars, however, molars are added posteriorly from the extension of the dental lamina (Juuri and Balic 2017). Mice and other rodents have a reduced number of teeth including 3 molars that are not replaced and continuously erupting incisors maintained with an active stem cell population in the cervical loops (Yu and Klein 2020). A major gap in knowledge about the dentition is evident since the majority of dental development has been characterized in the mouse model which does not undergo tooth replacement. Another reason for a lack of understanding of tooth replacement is that the actual process of tooth replacement occurs postnatally in humans (Ooë 1981) and all other diphyodont mammals such as pigs (Wang et al. 2014b), ferrets (Jarvinen et al. 2009) and shrews (Järvinen et al. 2008; Yamanaka et al. 2010). Thus far, only the molecular events of permanent tooth budding have been studied during the late fetal period in a small number human fetal specimens (Dong et al. 2014; Wang et al. 2014a) and other diphyodont mammals (Järvinen et al. 2008; Yamanaka et al. 2010). Animals that replace their teeth more than once are also being used in research studies. The bony fish, particularly those with oral teeth, can be followed in vivo. Indeed treating the fish with pathway inhibitors has uncovered some changes in tooth replacement and shape (Fraser et al. 2013). Many reptiles replace their teeth (crocodilians, snakes and lizards) throughout life. Previous longitudinal studies on alligators (Edmund 1962; Westergaard and Ferguson 1990; Westergaard and Ferguson 1987; Wu et al. 2013) and lizards (Brink et al. 2020; Cooper 1966; Grieco and Richman 2018; Kline and Cullum 1984) revealed waves of replacement that appear to be temporally and spatially regulated. Out of all these animals we have selected the leopard gecko, *Eublepharis macularius* for our research because the animals replace their teeth frequently in a highly patterned manner (Brink et al. 2022; Grieco and Richman 2018). Molecular studies have been carried out on the gecko, partially characterizing the main processes of tooth initiation and differentiation (Brink et al. 2021; Handrigan et al. 2010; Handrigan and Richman 2011; Kim et al. 2020; Zahradnicek et al. 2014).

In the leopard gecko, each functional tooth occupies a position at the margin of the jaws. There are approximately 40 teeth per quadrant of the mouth. Under each functional tooth is a tooth family attached to a long dental lamina. The 2^nd^ generation teeth consist of a mineralizing crown of enamel and dentin surrounding a dental papilla. The 3^rd^ generation teeth are detectable very early in development, at either bud or early cap stage. The enamel organs of geckos and other reptiles are similar to those in mammals consisting of an inner enamel epithelium that gives rise to ameloblasts, the intermediate stellate reticulum and the outer enamel epithelium. An important difference compared to mammals is that in the reptile there is no clear primary enamel knot, the main signaling centre in mammalian cap stage teeth. There is also not a clear stratum intermedium in reptilian enamel organs. The enamel organs are continuous with the dental lamina that connects the 3 generations of teeth together. The dental lamina continues past the 3^rd^ generation tooth to form the successional lamina. The functional teeth eventually are shed by a resorptive process and the next generation tooth erupts to fill the spot. The full tooth cycle from start to finish takes about 5 weeks in the gecko (Brink et al. 2022; Grieco and Richman 2018). This means that by harvesting gecko jaw tissues from any position we will capture all stages of tooth development.

Only 2 previous studies looked at the molecular basis of tooth shedding in reptiles (Fuenzalida et al. 1999; LeBlanc et al. 2023). These study employed Tartarate Acid Resistant Phosphatase staining (TRAP) to identify multinucleated, TRAP+ cells in *Liolaemus gravenhorsti* (lizard) and *Pantherophis guttatus* (snake). These TRAP+ cells were termed odontoclasts since they were located inside the pulp cavities. Similarly, human deciduous teeth have large numbers of TRAP+ odontoclasts that cause rapid resorption of the roots (Sahara 2001; Takada et al. 2004). To move the field forward it is necessary to profile resorbing teeth in a more comprehensive manner. Selecting the leopard gecko allows access to large numbers of teeth participating in physiological tooth resorption. In addition, collection of samples of non-resorbing developing teeth is feasible in the gecko. Both resorbing and developing teeth can be collected in a non-invasive manner (animals can return to the colony). These comparisons between shedding and developing teeth will also highlight developmentally regulated genes that may be novel in terms of a role in tooth formation. Conservation of some but not all genes with mammals is also likely. The reason we expect some differences is that geckos and non-crocodylian reptiles lack a periodontal ligament (Bertin et al. 2018). The periodontal ligament connects the root to the bone in mammals and is derived from the dental follicle(Nagata et al. 2022). Some snakes appear to have small remnants of Sharpey’s fibres that insert into the bone next to the teeth (LeBlanc et al. 2016; LeBlanc et al. 2017; LeBlanc et al. 2021) but these are radically reduced in these animals. Thus it will be interesting to see whether markers of the dental follicle are present in geckos and which cells express these genes.

Finally, the presence of stem cells has been best characterized in the mouse incisor (Chiba et al. 2020; Harada et al. 1999; Krivanek et al. 2020; Sharir et al. 2019). Using elegant lineage tracing and pulse-chase lineage tracing, it has been shown that the dental epithelial stem cells reside in the labial cervical loop (Krivanek et al. 2020; Sharir et al. 2019). We have also searched for putative stem cells in the gecko and found that the dental lamina is a source of stem cells (Brink et al. 2021; Handrigan et al. 2010). Markers for stem cells have only been identified to a very limited extent in the gecko (Handrigan et al. 2010). It is necessary to move towards single cell RNAseq and to enrich for epithelium to find more genes that identify rare populations of progenitor/stem cells. A previous study on bearded dragon used laser capture microdissection of the successional lamina to identify differences that are present in cells that will form successional teeth versus those that do not participate in tooth succession (Salomies et al. 2019). There were very few gene differences identified so a more comprehensive method is required with less disruption of the cells.

In this paper we used a combination of bulk and single cell, short read sequencing technology followed by validation with several methods. We report genes that were not previously associated with tooth development, tooth replacement or tooth resorption. In addition, markers of specific areas of mammalian teeth are found in gecko teeth, even though the homologous structures are not present. These unbiased studies will characterize the cell types within and around the teeth and will set the stage for future functional perturbations of physiological tooth replacement in the adult gecko jaw.

## RESULTS

Our aim was to take an unbiased approach to examine the differences in expression in functional teeth versus developing teeth. We predicted that the main processes that characterize developing teeth would include genes involved in odontogenesis, pulp cell maintenance and genes required to stimulate clastic cells differentiation. The scRNA-seq data was carried out with a different intent. Here we wanted to uncover cell populations that were recognizably dental but also expressed genes that have not been previously noted as a playing a role in tooth development. We also wanted to overcome the barriers of using a non-model organism, the leopard gecko in a full, unbiased transcriptome analysis.

### Quality control of samples used for bulk RNAseq

We first performed a principal component analysis of the samples using the count matrix from STAR aligner (Fig S2A). Principal component (PC) 1 explained 40.6% of the variability of the data, PC2 explained 22.47%. The functional tooth samples grouped together in the negative quadrants of PC1 axis while developing tooth samples grouped in the positive quadrants (Fig. S2A). Interestingly, PC2 accounted for the differences between animals (Fig. S2A). The samples belonging to animal 1 (two developing and 1 functional sample) were on the positive side of PC2 axis. The negative region of PC2 contained samples belonging to animal 2. Hierarchical clustering computed from the STAR matrix showed similarity within the functional tooth samples and within the developing samples (Fig. S1B).

A total of 16,954 genes (15086 with edgeR and 16306 with DESeq2) were detected across the samples. We performed a 5 pipeline analysis as described (Vuilleumier et al. 2019) to align and count the reads for each sample, generating 5 matrices. Two differential gene expression analysis were performed using edgeR and DESeq2 for each of the matrices. The false discovery rate (FDR) cutoff for significance was set at 0.05 for each individual pipeline, and since our procedure for combining the results of the 10 pipelines relies on the median ranking, a gene had to be called as significant in at least 5 of the 10 pipelines in order to make it to the final list (Table S1).

A total 1,139 transcripts were significantly differentially regulated after the 10 pipelines analysis (Table S1). Of these, 264 genes were downregulated and 875 were upregulated in developing teeth. This list of significantly regulated genes was further annotated using the NCBI tool BLASTx (Camacho et al. 2009) and the non-redundant database due to the high number of transcripts labelled as “unknown” or gene names without a clear homolog. We started with 34% of genes (390/1139) that fell into this category. After the blastx annotation, the list of unknown genes decreased to 19% (212/1139). These efforts were necessary in order to perform post-hoc analyses such as Gene Ontology to identify enrichment for certain biological processes. The reported p-value and log_2_ fold change in volcano plot were computed by applying edgeR to each of the kallisto matrix (Fig. 1A). The same applies for the DESeq2 (Fig S3A).

**Figure 1.**
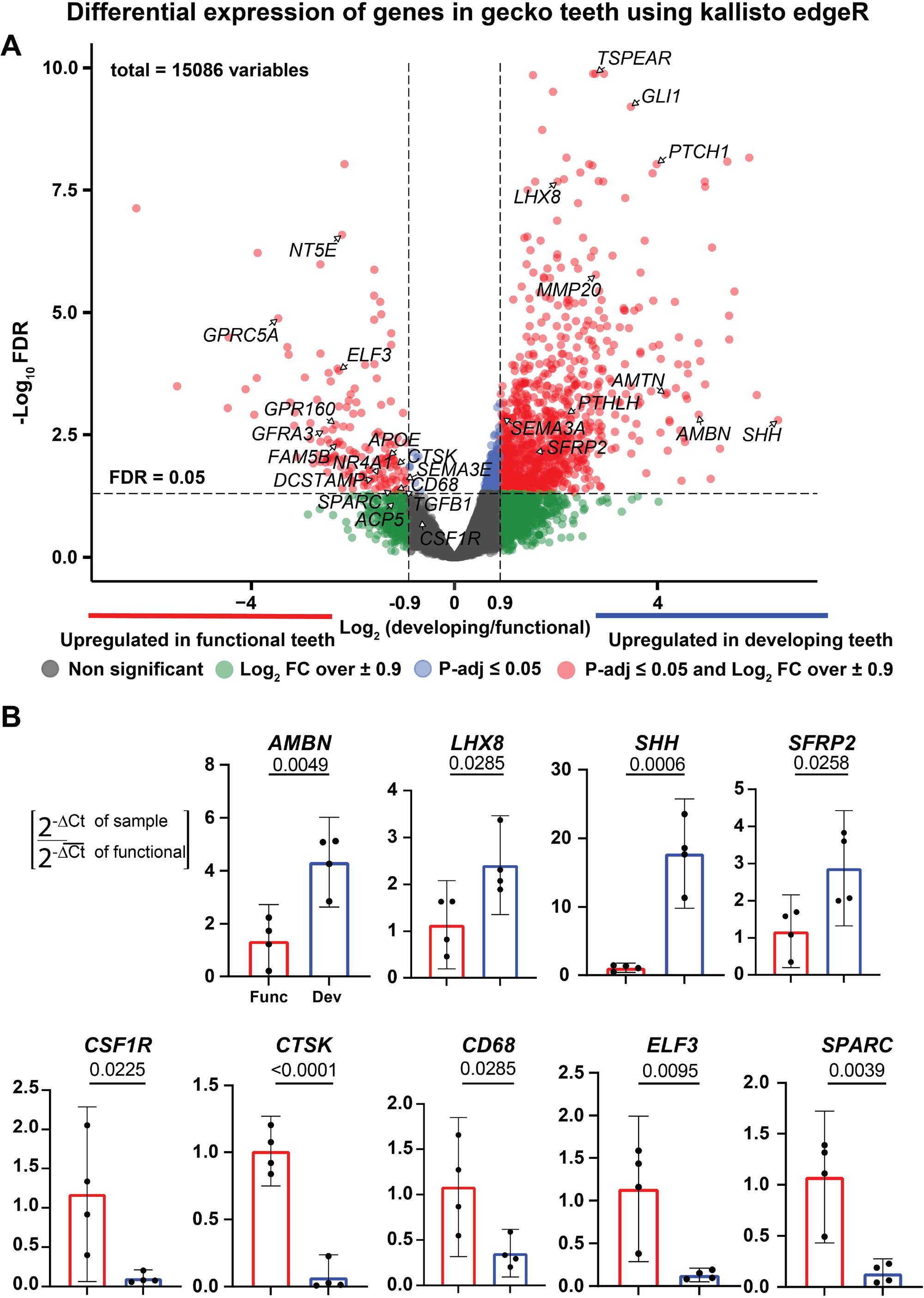
A) Volcano plot of transcripts sequenced in bulk RNAseq created with Kallisto and edgeR. Dots in red are transcripts with a p-value <0.05 and a Log_2_ of the ratio between the expression level of developing and functional teeth over ± 0.9. Transcripts traditionally associated with tooth development (*LHX8, AMBN, PTHLH, SFRP2* and *SHH*) are on the right side of the plot. Genes more associated with functional teeth are on the left side of the plot. Many of the genes have not been characterized in the tooth (*NT5E, ELF3, SEMA3A, SEMA3E, SPARC, FAM5B, GFRA3*). Others are expressed in cells with odontoclast/clastic activity (*CSF1R* and *ACP5*). *FST* does not appear in the edgeR analysis since this gene was below the cut-off. Fewer genes are plotted in edgeR than in DESeq2 (Fig. S3A). B) Independent samples were used for validation with qRT-PCR where the reference sample is always the functional teeth (95% confidence interval and the ratio of 2^-^Δ^CT^ values shown). Statistical significance for qRT-PCR was calculated using unpaired, two tail t-tests.

### Transcripts upregulated in developing teeth

Developing teeth collected during the microdissections are primarily in bell stage since we rely on calcein incorporation and this dye is only incorporated once mineralization has taken place (bell stage). As expected, genes required for early ameloblast differentiation were enriched in developing teeth. Ameloblastin (AMBN) is second most abundant proline-rich matrix glycoprotein in enamel (Moradian-Oldak and George 2021). *AMBN* was significantly upregulated in developing teeth (log2FC = 4.66; 5-fold enriched; Fig. 1A,B; S3A). Amelotin (*AMTN*) and *MMP20* (Table S1; Fig. 1A, S3A) also function during amelogenesis (Bartlett and Simmer 2015; Hu and Simmer 2007).

In the present study the bulk dissection of bell stage teeth captured inner enamel epithelial transcripts. *Shh* (Sonic Hedgehog) is expressed in the primary enamel knot and inner enamel epithelium in placode, bud, cap and bell stage of mouse teeth (Vaahtokari et al. 1996). We have also published expression of *SHH* in gecko teeth, particularly in the inner enamel epithelium of bell stage teeth, prior to differentiation of the ameloblasts (Handrigan and Richman 2010). *SHH* was more highly expressed in developing teeth (Fig. 1A; log2FC = 6.19) and this was validated with qRT-PCR (Fig. 1B; 18-fold higher than functional teeth). Target genes of the hedgehog (Hh) pathway are also upregulated in developing teeth including *PTCH1* (log2FC = 3.8; Fig. 1A, S3A) and *GLI1* (log2FC = 3.36; Fig. 1A, S3A). Another dental epithelial gene, *EDAR* (Ectodysplasin receptor, log2FC = 2.09; Table S1; Fig. 1A, S3A) is also more highly expressed in developing teeth. In the mouse, *Edar* is exclusively expressed in the dental epithelium at placode, bud and cap stages (Laurikkala et al. 2001). *Edar* is also strongly expressed in the primary enamel knot of mice (Laurikkala et al. 2001) however gecko teeth do not appear to have a primary enamel knot (Handrigan and Richman 2011). Thus the function of EDAR in the replacing reptile dentition may be different than in mammals. *TSPEAR* (Thrombospondin type laminin G domain and EAR repeats) was also upregulated in developing teeth (log2FC =2.59; Table S1; Fig. 1A, S3A). Mutations in human *TSPEAR* produce ectodermal dysplasia type 14 (OMIM: 618180), which is also characterized by hypodontia (Peled et al. 2016).

We identified several genes that are expressed more highly in the dental mesenchyme. For instance, *LHX8* is a transcription factor that is expressed in early neural-crest derived mesenchyme and in dental mesenchyme of developing mouse molars (Grigoriou et al. 1998; Zhao et al. 1999; Zhou et al. 2015) (log2FC=1.82; Fig. 1A,B; Fig. S3A). Follistatin (*FST*) is highly expressed in dental mesenchyme in mouse teeth (Wang et al. 2004) and is also increased in developing gecko teeth (log2FC = 7.33; Table S1; Fig. 1A, below cut-off for DESeq2). Interestingly, *Sfrp2* (Secreted Frizzled Related Protein 2) a WNT (Wingless-related protein) antagonist) is expressed in the dental follicle in the mouse dentition (Sarkar and Sharpe 1999) (Krivanek et al. 2020)and in gecko developing teeth (log_2_ = 1.41; Fig. 1A,B; Fig. S3A). Overall, the expression of genes in developing gecko teeth contains genes found in the embryonic mouse molar but there is a subset of genes such as *EDAR* and *SFRP2* that may have radically different spatial locations of transcripts compared to the mammal.

### Transcripts upregulated in functional teeth

We were also interested in identifying genes relevant to tooth homeostasis by profiling expression in functional teeth. *SPARC* (Secreted acidic cysteine rich glycoprotein or Osteonectin), which has an important role in biomineralization of bone, was differentially expressed in functional teeth (log2FC = -1.4; Fig. 1A,B; S3A). Mutations in this gene in human cause Osteogenesis Imperfecta type XVII (OMIM# 616507). Thus far, tooth phenotypes have not been associated with mutations of the *SPARC* gene (Mendoza-Londono et al. 2015). In addition, we identified two novel genes not previously known to be involved in tooth function including *ELF3* (log2FC = -2.43; Fig. 1A,B; S3A) and *NT5E* (log2FC = -2.43; Fig. 1A,B; S3A). *Elf3* (E74-like factor 3), is expressed in salivary gland (Song et al. 2022), hair follicle (Choi et al. 2008) and mammary gland (Choi et al. 2009). Since all of these are ectodermal derivatives, it is possible that expression of *ELF3* in functional gecko teeth is in the only areas with residual epithelium, the cervical loops. There are no mouse studies that examined expression of *Elf3* in the dentition. *NT5E* is a nucleotidase that when mutated causes ectopic calcification of joints and arteries (OMIM: 211800) (St Hilaire et al. 2011). Again, *Nt5e* has not been observed to play a role in the mouse dentition. We also identified significantly enriched transcripts for *GPRC5A* in functional teeth (G protein-coupled receptor, family C, group 5, member A; log2FC=-3.74; Fig. 1A, S3A). *GPRC5A* is a 7-transmembrane G protein coupled receptor, whose expression is induced by retinoic acid (Cheng and Lotan 1998). The ligand for GPRC5A has not been identified (Richter et al. 2020) and there is no known role for this gene in odontogenesis.

### Bulk RNAseq identifies genes associated with root resorption in functional teeth

Interestingly, genes expressed by osteoclasts such as *ACP5* (Fig. S3A) and *CSF1R* (Fig. 1A, B; S3A), were upregulated in functional teeth. In addition, *PTHLH*, a potent inducer of RANKL (Rao et al. 2018) a classic osteoclast differentiation factor, was present in developing and functional teeth (Fig. 1A, S3B). The macrophage marker *CD-68* (log2FC _=_ -1.34) is upregulated in functional teeth (Fig 1A,B; S3A). *CD68* is expressed in fully differentiated osteoclasts and not their precursors (Jackson et al. 2017). *TGFB1* signals via the transcription factor Smad4 and there is evidence that TGFB1 protein activates ostoclastogenesis in mice (Morita et al. 2016) and was differentially expressed in functional teeth(Fig. S3A,B; log2FC = - 1.17). The location of all of the aforementioned gene transcripts and their role in the adult gecko tooth has not been explored.

We next examined the functional teeth in detail to see whether there were odontoclasts within the pulp cavity. The odontoclast is similar to an osteoclast but has not been as well characterized (Wang and McCauley 2011). As a proof of principal we examined the gecko egg tooth for presence of odontoclasts (Fig. S4A). The egg tooth is rapidly shed at the time of hatching (Fons et al. 2020). The TRAP stain (Tartrate resistant acid phosphatase is coded by the *ACP5* gene) is a marker for odontoclasts (Fuenzalida et al. 1999; Lee et al. 2020; Sahara et al. 1996; Takada et al. 2004). Indeed, TRAP+ cells are inside the egg tooth (Fig. S4B). The functional teeth showed the presence of many multinucleated cells that are TRAP+ inside resorption pits (Fig. 2A,A’,B,B’’, S4C,E,F,G,H,I). The patterns of invasion into functional teeth (approximately every other tooth (Fig. S4H,I). There were no multinucleated cells in developing teeth (Fig. 2A’’,B’’’; S4F,I).

**Figure 2.**
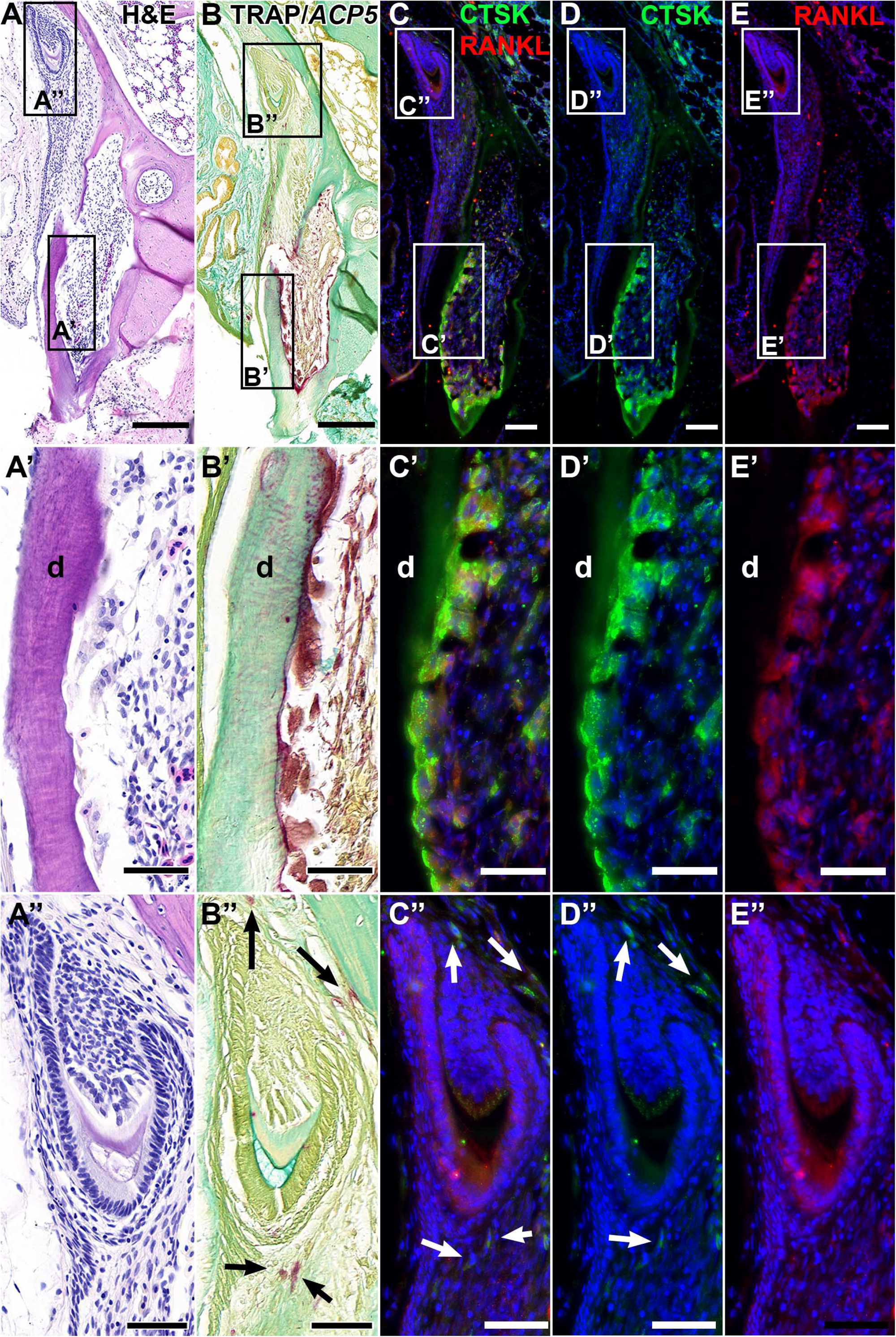
Near adjacent sections stained for H and E (A-A’’), TRAP staining (B-B’’) and double immunofluorescence staining with antibodies to Cathepsin K and RANKL (C-E’’). A) A tooth family with a a functional tooth (A’) and a developing tooth (A’’). B-B’’) TRAP positive, multinuclear odontoclasts are attached to the inner pulpal surface. B’’) Developing teeth do not have TRAP+ cells but there are positive cells next to the bone surface (arrows). C,C’) Adjacent sections show strong staining for CTSK and RANKL overlapping areas of TRAP+ cells. C’’) Developing teeth have RANKL staining in the ameloblast layer but no CTSK (D’’). There are cells next to the tooth bud and the bone that are TRAP+ cells in one section (B’’) and CTSK positive in adjacent sections (C’’, D’’). E’’) There does not appear to RANKL expression in the cells next to the tooth that are TRAP+. Scale bars: A-B, 200 µm; C-E, 100 µm; A’-E’ 50 µm: A’’-E’’, 50 µm.

There are other markers that identify earlier stages of osteoclast formation and these may also be expressed in odontoclasts (Dai et al. 2020; Yasuda 2021). We used antibodies that recognize RANKL (Rank Ligand) and CTSK (Cathepsin K). The multinucleated cells that stained with TRAP were also positive for CTSK and RANKL (Fig. 2C’-E’). Developing teeth did not have positive cells in the pulp cavities (Fig. 2C’’-E’’). However, surrounding the developing teeth (between the teeth and the bone) there were small groups of TRAP+ and CTSK+ cells that were negative for RANKL (Fig. 2B’’-E’’). These cells are also mononuclear and therefore could be precursor monocytes that are migrating towards the pulps of functional teeth.

The robust expression of the CTSK antibody prompted us to search the RNAseq data for the transcripts. Unfortunately, the CTSK gene has not been annotated in the draft genome (Xiong et al. 2016). We therefore re-annotated the genome de novo using our RNAseq data as hints and successfully identified the CTSK gene (see Methods for details). We performed the transcriptome analysis again and found significantly elevated *CTSK* in gecko functional teeth (Fig. 1A,B; S3A). This one gene exemplifies some of the issues with the draft *E. macularius* genome(Xiong et al. 2016).

An unexpected class of genes that was expressed in gecko functional and developing teeth belonged to a class of chemotaxis molecules, the Semaphorins. *SEMA3A* and *SEMA3E* are primarily known as guidance molecules for the nervous system and angiogenesis. *SEMA3E* was downregulated in developing teeth when compared to functional teeth, whereas *SEMA3A* had the opposite trend. *SEMA3E* differential expression was significant in both edgeR and DESeq2 (Fig. 1A; S3A), while *SEMA3A* was significant only in DESeq2 (Fig. S3A). SEMA3A has been shown to be a chemo repellant that controls timing of tooth innervation in the mouse (Luukko and Kettunen 2016; Moe et al. 2012a; Moe et al. 2012b; Shrestha et al. 2014). SEMA3E is best known for regulating vascular patterning in mammals and not odontogenesis (Kim et al. 2011). Moreover, does not appear to be *Sema3e* expressed in developing mouse teeth (Fu et al. 2017).

We localized transcripts for both *SEMA3A* and *3E* in the gecko dentition using RNAscope (Fig. 3A-D’). *SEMA3E* was expressed in the odontoblasts and in mesenchyme external to the tooth root (Fig. 3A-A’’). In developing teeth, *SEMA3E* was detected in the odontoblast layer (Fig. 3B,B’). *SEMA3A* was expressed in the differentiated odontoblasts of functional teeth and external to the teeth (Fig. 3C’C’’). In developing teeth, *SEMA3A* was mostly expressed in the apical papilla in between the cervical loops, a pattern not seen with *SEMA3E* (Fig. 3D,D’). There was also expression of *SEMA3A* external to the developing tooth (Fig. 3D’). It is interesting that *SEMA3A* acts as a chemorepellant for nerves in the murine dentition (Luukko and Kettunen 2016). We suspect that the strong apical presence of *SEMA3A* delays nerve ingrowth until the tooth is more fully developed. There may also be novel functions of SEMA3A that will need to be explored in functional experiments.

**Figure 3.**
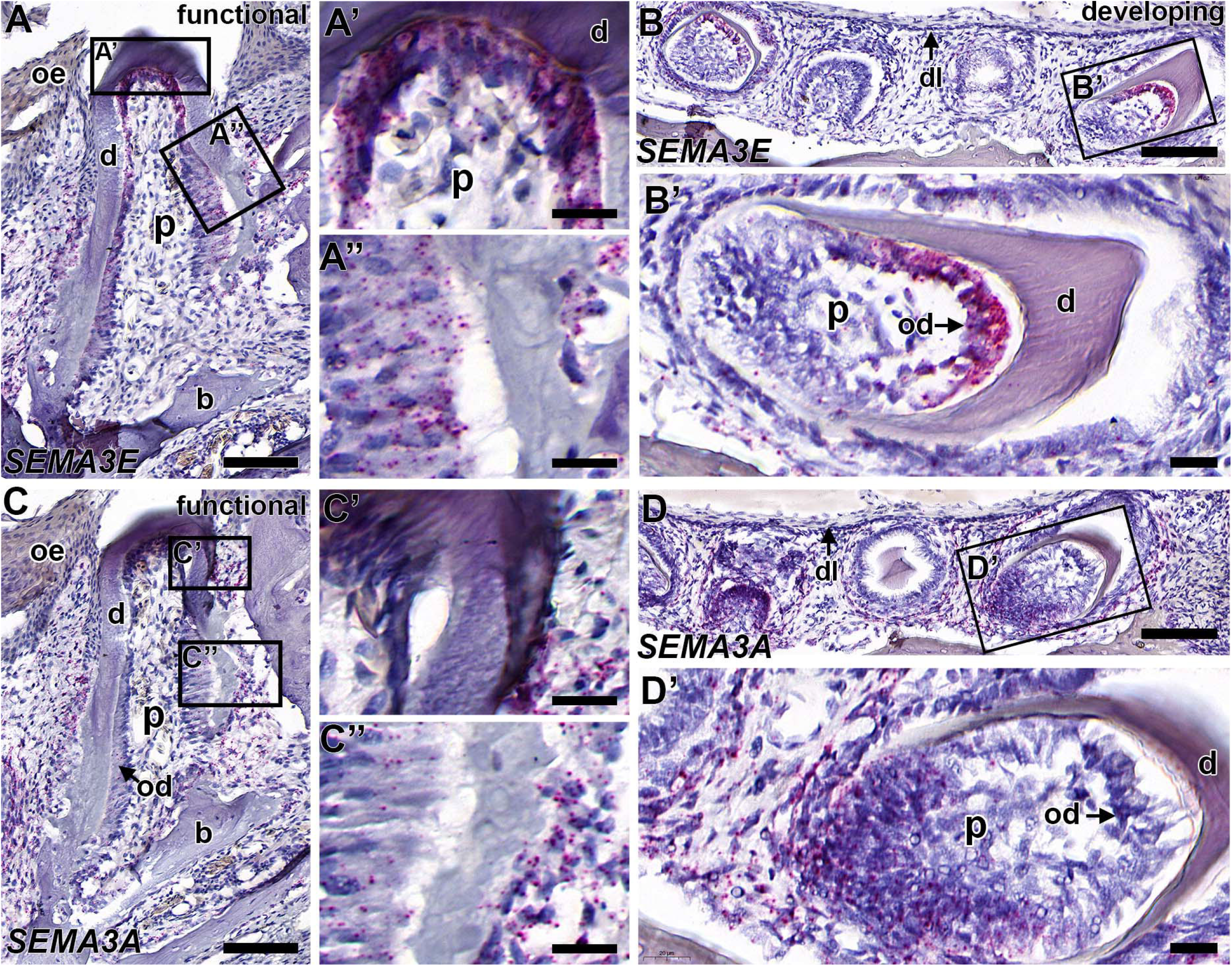
RNAscope in situ hybridization with probes that recognize *SEMA3E* (A and B) or *SEMA3A* (C and D). Near-adjacent mandibular sections of functional (A and C) and developing teeth (B and D). The expression of *SEMA3E* is in the mature odontoblasts of functional teeth (A-A’’) while *SEMA3A* is most abundant in odontoblasts actively secreting dentin (C-C’). In developing teeth *SEM3A* is expressed in the newly differentiated odontoblasts (B’’). *SEMA3A* is expressed in the apical dental papilla (D,D’). Key: b-bone, d-dentin, dl-dental lamina, oe -oral epithelium, od-odontoblast layer, p-pulp. Scale bars: A, B, C and D = 100 µm; A’, A’’, C’, C’’, B’ and D’ = 20 µm.

### Single cell RNAseq identifies clusters of dental epithelial and mesenchymal cells

The reasons for carrying out bulk RNAseq specifically on the dissected teeth were that we could be sure that all genes identified would be specific to the teeth and not surrounding tissues (Fig. S1A). The comparison between bell stage and functional teeth was designed to identify genes required for tooth shedding, eruption and those needed for crown completion. Other cells adjacent to the developing teeth, particularly the dental lamina and successional lamina are critical for tooth initiation in the adult gecko (Brink et al. 2021). We turned to single cell RNAseq in order to isolate cells comprising the tooth field deep in the jaws. This dissection included dental lamina, successional lamina, early tooth buds, bell stage teeth, oral connective tissue and mucosa (Fig. S1B). We obtained 6,197 cells, 68,380 post normalization mean reads per cells and 492 median genes per cell (Fig. 4A; S5).

**Figure 4.**
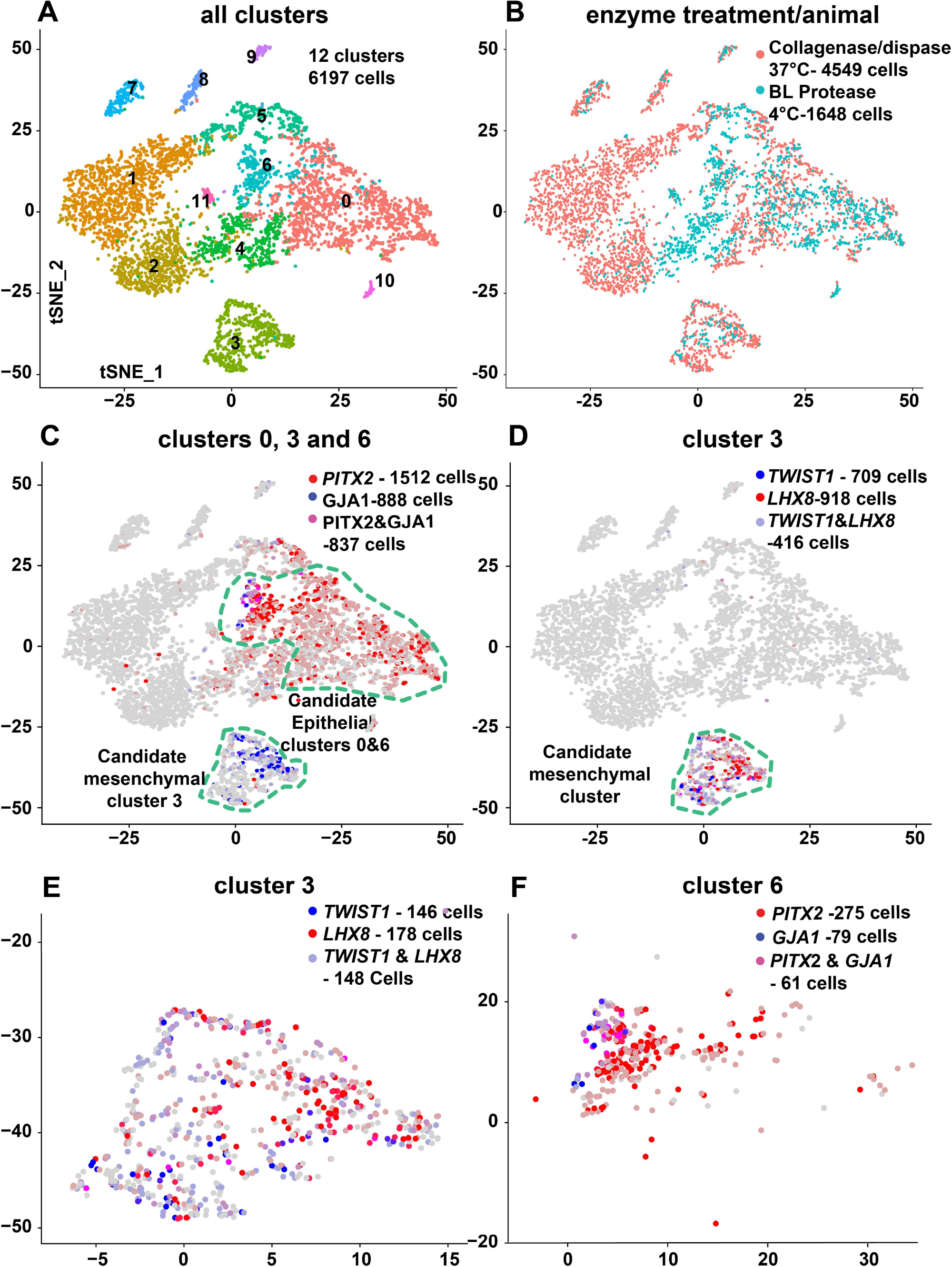
T-distributed stochastic neighbour embedding (tSNE) plot of single cell RNA sequence. A) The 6197 cells were put into 12 distinct clusters. B) Cells colored according to enzymatic treatment used to dissociated the cells from the tissue. The cells were derived from different geckos. C) Putative epithelial clusters 0,3,6 encircled overlaid with the level of expression of two dental transcripts (*PITX2* and *GJA1*). A putative dental mesenchymal cluster showing the levels of expression of a dental mesenchyme marker (*GJA1*). D,E) Cluster 3 showing the expression of *TWIST1* and *LHX8*. F) Cluster 6 contains cells from the dental epithelium as shown by high expression of *GJA1* and *PITX2*. Number of cells with counts over 1 are indicated in each panel.

We carried out two different cell isolation protocols, the typical Collagenase/Dispase dissociation mixture at 37°C and a cold temperature protocol (Adam et al. 2017)(Fig. 1B). The body temperature of the gecko is 28-32°C so it was necessary to try a different method that may be more compatible with ectothermic animals. The warm temperature isolation released more cells than cold temperatures (Fig. S5A,D,G compared to S5B,C,E,F,G). The quality of reads was higher at cold temperatures (Fig. S5C,F). Using the Cell Ranger pipeline, the number of genes per cells and unique molecular IDs or bar codes were also higher in the *B. lichenoformis* method (Fig. S5G).

Next, we began by looking for patterns of cells isolated with the two different isolation methods from two different animals (Fig. 4B). Using the Seurat package in R, we computed 12 clusters and a list of differentially expressed transcripts (Table S3). We used FindClusters (resolution = 0.2). All cells were included with nFeature or gene IDs between 175 and 2600. As with the bulk RNAseq we used the Blastx (Camacho et al. 2009) function to identify the human orthologous gene name for the 45% unknown genes (662 genes out of 1458 unknown genes). After performing Blastx, the number of non-useful annotations decreased to 21% (307/1458). A gene ontology analysis was carried out using GO Consortium and we used the Biological processes (Table S4) to assign identities to 6 out of 12 clusters, 2 with certain dental identity (clusters 3 and 6) and one with a mix of dental and non-dental cell types (0). There were other clusters containing non-dental cell types (1,5,9) (Fig 4A).

Most of the clusters were comprised of a mix of cells isolated with the cold and warm enzyme treatments, however cells in clusters 1,2,3,5,7,8 and 9 originated almost exclusively from the animal (maxilla) dissociated with collagenase/dispase (Fig. 4B; S6A,B). Clusters 4,6,10 and 11 are mainly associated with cells processed by the cold temperature isolation (animal 2, mandible). Cluster 0 also had 40% of the cells originated from the cold technique. Thus both methods have advantages and disadvantages and combining the two types of isolation methods helps to ensure populations of cells are not missed. It is more likely that the type of enzyme was responsible for the differences in cells that were isolated rather than the animal or the region of the jaw. We have seen interchangeable results in the maxillary or mandibular teeth in previous work (Brink et al. 2021) and data not shown.

### Genes not annotated hinder cluster identity assignment

Several clusters had small numbers of differentially expressed genes and using the GO biological process enrichment, identities were not assigned (Table S3; Fig. S11D-O). We identified cluster 1 as most likely being composed of T-cells (Fig. S9B-F, M,N). Key markers of T-cells include *TESPA1*, which is involved in T-cell development (Wang et al. 2012). In addition, the HLA group is mainly restricted to cluster 1 (Fig. S9F). Cluster 2 is most likely composed of lymphocytes and other white blood cells (Fig. S9G-I). In cluster 4 we identified enrichment of *TMEM249* and *C3* (Fig. S9M,N) a gene coding for complement which is in the innate immunity cascade). Cluster 5 contained proliferative cells of unknown identity. The cell express markers associated with cell proliferation like *PCNA, CDK1* (Gautier et al. 1988) and the PCNA associated factor *PCLAF* (Emanuele et al. 2011) (Fig. S10B,C). Clusters 7 and 8 contain cells secreting cytokines and complement factors (Fig. S10D-I) but the exact cell type is unclear. Cluster 9 expressed genes associated with angiogenesis (Fig S10J-L) like *ROBO4* which is highly expressed in endothelial cells (Jones et al. 2008). Cluster 10 (Fig. S10M-O) and cluster 11 (Fig. S11B-D) have no clear identity.

### Clusters with epithelial cell identity

Cluster 0 was confirmed as epithelial since multiple keratin transcripts were present (Table S3, Fig. 4C; Fig S8B-F; *KRT 4, 5, 12, 14, 23, 24*). In humans, keratin 5 (a type I keratin) and 14 (a type II keratin) form a pair that is expressed in the keratinocytes of the basal layer of the epidermis, the basal cells of endodermally-derived glands and other stratified epithelia (Porter and Lane 2003). Mutations in *KRT5* or *KRT14* produce epidermolysis bullosa simplex (Porter and Lane 2003). Antibodies to KRT14 strongly stained cells of the human dental lamina (Domingues et al. 2000). Cluster 0 also contains cells that express a known dental identity gene, *PITX2* (Fig. 4C)(Thesleff 2014) and *SOX2* a, dental epithelial stem cell marker (Fig. S8K)(Yu and Klein 2020). Mutations in *PITX2* cause Axenfeld Rieger syndrome type I (OMIM: #180500), which also features dental anomalies. The gap junction gene, *GJA1* (codes for Connexin 43) is also expressed in the epithelial cluster 0 (Fig. 4C). Mutations in *GJA1* cause oculo-dento-digital dysplasia syndrome (OMIM: #164200). Therefore, cluster 0 consists of dental and non-dental epithelial cells.

Cluster 6 had a higher proportion of dental epithelial gene transcripts (Table S3; Fig. 4F; Fig. 5A). Cluster 6 expressed *PITX2*, which is strongly expressed in the enamel organ of developing teeth and successional lamina (Fig. 5A,B). We previously mapped in detail the expression of PITX2 in the leopard gecko dentition (Brink et al. 2021). *GJA1* which codes for Connexin43 is also expressed strongly in cluster 6 (Fig. 5A). CX43 protein is mainly restricted to the enamel organ of developing teeth (Fig. 5C) but absent in the dental and successional laminae. Ameloblasts in the inner enamel epithelium at secretory stages have intense signal particularly in the lateral aspect of the cell membranes (Fig. 5C, S7B-B’’) as described in the mouse (Toth et al. 2010). Mice with a heterozygous loss-of-function mutation in *Gja1* have poor quality, thin enamel (Toth et al. 2010). We verified that CX43 protein is present at lower levels in the odontoblasts and dental papilla (Fig. 5C’, S7B’). Thus, in the gecko, CX43 was not exclusively expressed in epithelial cells, as reported in the mouse (Toth et al. 2010). Cells in cluster 6 express other genes found in the enamel organ including *SHH* (Table S3). *SHH* was previously shown to be expressed in the embryonic leopard gecko enamel organ (Handrigan and Richman 2010). Cluster 6 cells expressed *Fermt1* (Rostampour et al. 2019), *Irx1*(Yu et al. 2017), *Ednrb* (Neuhaus and Byers 2007) and *Itgb4* (Salmivirta et al. 1996) all of which are expressed in the mouse enamel organ. Markers of fully functional ameloblasts were not identified in cluster 6, even though these genes were found in the bulk-RNA-seq data for developing teeth. The reason may lie in the difference in dissection methods used for scRNA-seq. In this enzyme protocol we did not use calcein labeling to identify mineralizing teeth. All such teeth would have been filtered out during the cell isolation process.

**Figure 5.**
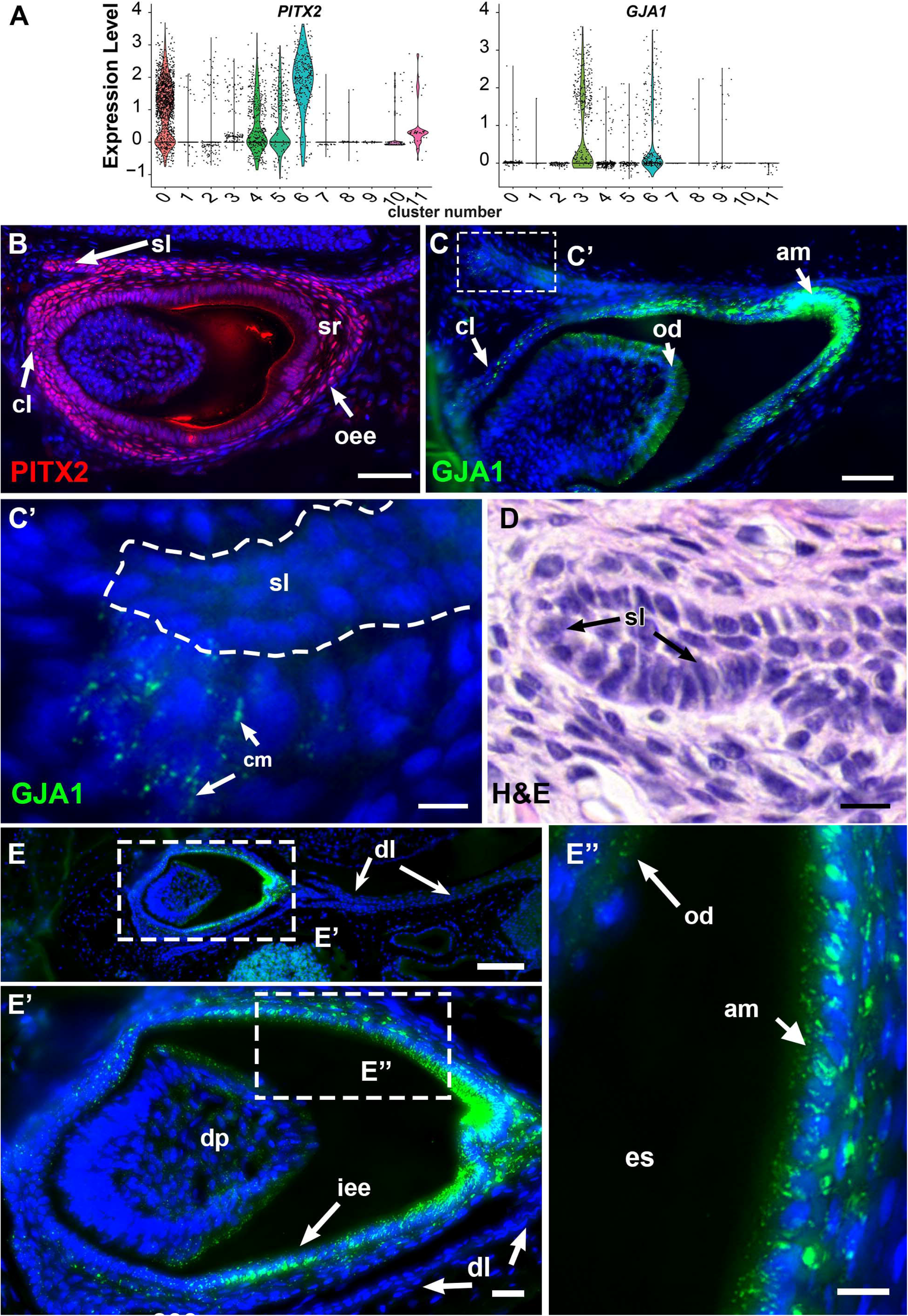
Spatial localization of proteins coded by 2 genes more highly expressed in developing gecko teeth. A) Violin plot of the scRNA-seq clusters showing the expression levels for *PITX2* and *GJA1*. B) Immunofluorescence staining with PITX2 antibody showing expression in all layers of the enamel organ and successional lamina. C) Expression of GJA1 protein in the differentiating ameloblasts and a few cells in the cervical loops. C’) At high magnification cells in the dental papilla of the newest successional tooth express GJA1. C’’’) Near-adjacent section of C’ stained with H&E showing the structure of the successional lamina and mesenchyme. E) GJA1 immunofluorescence staining of developing tooth but no expression in the dental lamina. E’) Expression in the ameloblasts of the inner enamel epithelium as well as a small amount in the odontoblasts of the dental papilla. Key: am – ameloblasts, cl-cervical loop, cm-condensing mesenchyme, dl-dental lamina, dp-dental papilla,es-enamel space, iee-inner enamel epithelium, od-odontoblast, sl-successional lamina, sr-stellate reticulum. Scale bars: Bars in B,C = 50 µm; bars in C’,D,E’’ = 10 µm; E=100 µm; E’=20 µm

### Cluster 3 has dental mesenchyme identity

Of all the cell groups, cluster 3 most closely resembled dental mesenchyme (Fig. 4D,E). As predicted by the immunostaining, we did find expression of *GJA1* in cluster 3 cells (Fig. 4D). Markers such as *LHX8, TWIST1, TGFB1, ALX1, SFRP2* and *OSR2* (Fig 5DE, Fig S9J-L) were highly differentially expressed compared to other clusters (Meng et al. 2015; Zhou et al. 2015). TWIST1 protein is expressed strongly in mesenchyme and in particular in dental mesenchyme of the mouse (Higashihori et al. 2017). *Lhx8* (formerly known as *Lhx7*) is strongly expressed in craniofacial mesenchyme including the tooth bud mesenchyme (Grigoriou et al. 1998) as well as in the root mesenchyme. The *Osr2* transcription factor is expressed in the mesenchyme lingual to the dental lamina in both the maxilla and mandible of mice at stage 11.5. Deletion of *Osr2* in the mouse causes molar duplication (Zhang et al. 2009). Cluster 3 also expressed *COL1A1* (Fig. S11A; Table S2) which is widely expressed in all connective tissues and in the dental papilla of developing teeth (Fig. S11B-B’’’’). There is no expression of *COL1A1* in the epithelium (Fig. SB’’’’). Cluster 3 does not express *SEMA3A*. In mammals the tissue that surrounds developing teeth is a sac-like structure called dental follicle that expresses IGFBP5 and ACTA2 (Hermans et al. 2022). Here we found IGFBP5 expressed in cluster 3 (Fig. S12A) but very few cells expressed ACTA2 (Fig. S12C). Therefore, cluster 3 may include cells from a dental follicle-like structure. Other markers of dental follicle identified by others include *Bmp3, Spon1, Hhip, Aldh1a2, Gdf10, Foxc2, Cnn1, Ntn1, Pdzrn4, Ltbp2, Hpse2, Podnl1* (Krivanek et al. 2020). At least as far as we know based on the annotations of the gecko genome, none of these genes were significantly enriched in the gecko scRNA-seq.

### Localization of *PTHLH* and *SFRP2* transcripts in and around developing teeth

*PTHLH* (Parathyroid hormone-like hormone), was present in the developing and functional teeth in bulk RNAseq (Fig. 2A, B) and was also statistically significantly enriched in the dental epithelial cells of cluster 6 (Fig. 6A, Table S3). PTH signaling is required for tooth eruption in mice (Philbrick et al. 1998; Takahashi et al. 2019) and humans (Frazier-Bowers et al. 2014; Frazier-Bowers and Vora 2017). Mouse *Pthlh* is expressed in the outer enamel epithelium and stellate reticulum (Philbrick et al. 1998). The same group reported the expression of *Pthlh* in the stellate reticulum of 8 day postnatal rats (Nakchbandi et al. 2000). The lineage tracing experiments yielded signal in the dental follicle using Pthlh-mCherry knock-in mice (Takahashi et al. 2019). There are also scRNA-seq data with conflicting data concerning the epithelial expression of *Pthlh*. In another study using RNAscope, Pthlh transcripts in the incisor cervical loop (Sharir et al. 2019). Similar to the original mouse in situ data, we found that gecko *PTHLH* was mainly located in the dental epithelium. *PTHLH* probe was localized to cells of the inner enamel epithelium (Fig. 6A’,C), stellate reticulum (Fig. 6A’’,E), successional lamina (Fig. 6D) and secretory ameloblasts (Fig. 6E). There were a few transcripts present in the dental papilla (Fig. 6C). We also examined the odontoclasts to determine whether these cells expressed *PTHLH* RNA. However, there was no expression. It will be interesting to see where the receptor for PTHLH - PTH1R – is expressed. Then we will understand the cell target of the epithelial-derived PTHLH. The most abundant expression of *PTHLH* was in the stellate reticulum of second generation teeth, next in line to erupt into the mouth (Fig. 6A-A’’). Therefore, the expression of *PTHLH* in the gecko is more similar to those mouse studies that reported mainly enamel organ expression (Philbrick et al. 1998; Sharir et al. 2019).

**Figure 6.**
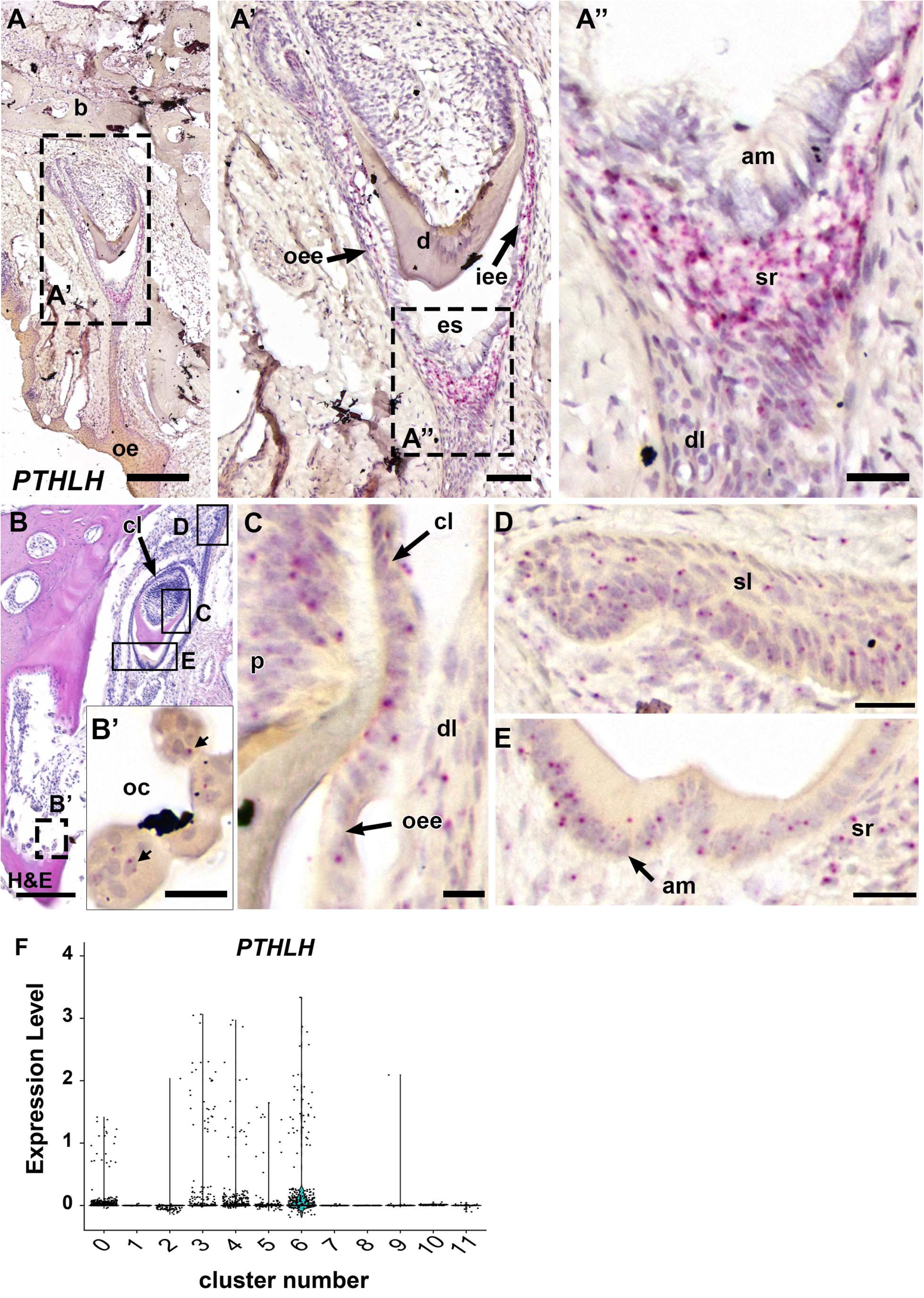
Expression of *PTHLH* RNA in gecko teeth. A) A 2^nd^ generation tooth beginning to resorb functional tooth. A’,A’) The stellate reticulum expresses signal between the inner and outer enamel epithelium. The dental lamina and fully differentiated ameloblasts do not express *PTHLH*. B) A resorbing functional tooth, a second generation tooth and the successional lamina. B’) There is very little signal within the multinucleated odontoclasts. C) Expression is present in the outer enamel epithelium as it extends to form the cervical loop. D) There is light signal in the successional lamina. E) Signal is present in secretory ameloblasts and the stellate reticulum. F) Violin plot showing distribution of *PTHLH* expressing cells in the clusters. The most abundant signal is in cluster 6, the dental epithelial cluster. Key: am-ameloblast, cl-cervical loop, cm-condensing mesenchyme, iee-inner enamel epithelium, oc-odontoclast, oee-outer enamel epithelium, sl-successional lamina, sr-stellate reticulum. Scale bars: A,B=200µm, A’,A’’=50 µm; B’,D,E=20 µm, C=10 µm

The expression of the WNT pathway antagonist, *SFRP2*, was significantly higher in developing teeth and was specifically enriched in the mesenchymal cluster 3 in scRNA-seq data (Fig. S12B). *SFRP2* is a marker of the apical papilla in humans (Krivanek et al. 2020). *Sfrp2* is expressed around the mouse molar (Sarkar and Sharpe 1999) and incisor dental follicle (Krivanek et al. 2020). We were interested to examine the gecko dentition, where it is difficult to discern a dental follicle by histological stains (Fig. 7A,C,E,G). There was signal for *SFRP2* adjacent to the developing teeth. The transcripts were localized to a band of mesenchymal cells surrounding developing bell stage teeth (Fig. 7A,B). In addition, there is expression around the successional lamina (Fig. 7C-F). The bud stage teeth are also surrounded by *SFRP2* expressing cells (Fig. 7G,H). There appears to be expression joining the developing tooth to the functional tooth (data not shown). We do not see expression of *SFRP2* in the dental epithelium. The function of this very defined, SFRP2+ population in the replacing gecko dentition requires further investigation.

**Figure 7.**
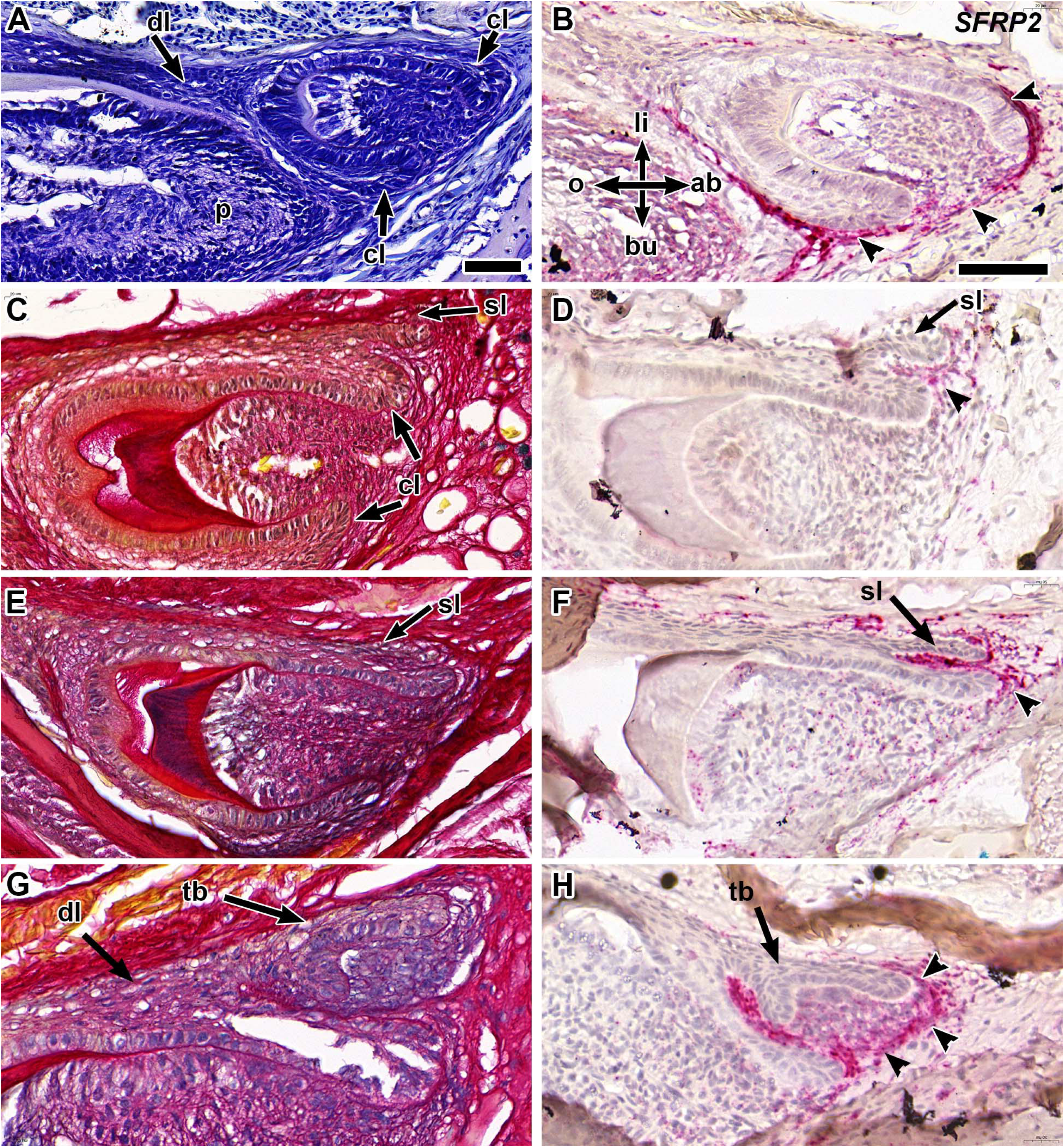
RNAscope in situ hybridization with a probe that hybridizes to *SFRP2*. Transverse maxillary sections stained with toluidine blue (A) or Picrosirius red (C,E,G). Near adjacent sections hybridized to *SFRP2* probe and counterstained with hematoxylin (B,D,F,H). A) Third generation tooth bud attached to the dental lamina. B) Strong signal for *SFRP2* is localized to a band of mesenchymal cells surrounding the tooth bud (arrowheads). There is no signal in the dental papilla, cervical loops or the enamel organ. C) A second generation tooth bud in late bell stage with differentiating enamel and dentin. The successional lamina is budding off the lingual side. D) the SFRP2 signal is surrounding the successional lamina and adjacent to the cervical loop on the lingual side (arrowheads). E) 2^nd^ generation tooth in late bell stage with successional lamina budding from the lingual side. F) strong *SFRP2* signal next to the successional lamina (arrowheads). G) Bud-stage 3^rd^ generation tooth extending from the dental lamina. H) *SFRP2* signal is surrounding but not within the tooth bud (arrowheads). Key: ab-aboral, bu-buccal, cl-cervical loop, dl-dental lamina, li-lingual, o-oral, p – papilla, sl-successional lamina, tb – tooth bud. Scale bars: Bar in A = 50 µm and applies to A,C,E. Bar in B= 50 µm and applies to B,D,F,G,H.

## DISCUSSION

In this study we used unbiased transcriptome profiling of the healthy adult reptile dentition in the midst of physiological tooth replacement. All stages of the tooth replacement cycle were captured because we used two complementary methods of RNA collection. The bulk RNA-seq included functional teeth that participate in the normal ingestion of food. We also included functional teeth at the end of the life cycle. These teeth no longer have a functioning pulp and instead are invaded by odontoclasts. The odontoclasts rapidly resorb the dentin and the bone-dentin interface allowing teeth to be shed. The other half of the tooth replacement cycle was captured with single cell dissociation of the soft tissues in the jaws. The deepest, most aboral region always contains the nascent tooth forming field. By dissecting jaw regions across several tooth families, we included a range of tooth initiation stages from the successional lamina, tooth bud, cap and bell stage. The dental lamina which is continuous around the jaw was also dissociated and included in the samples. Indeed, using two different enzymatic treatments, helped to improve the yield of cells in the dental epithelial and dental mesenchyme clusters. The data has revealed both conservation and novel aspects of odontogenesis not present in the mouse model.

### Cold versus warm isolation methods in scRNA-seq preferentially enrich for different cell types

The novel approach of using scRNA-seq on a reptile motivated us to test two different methods of cell isolation. We used a warm method that requires 37° C temperatures even though this is above body temperature of lizards and most ectothermic reptiles (28-29°C). Thus far we are the only group to use *B. lichenoformis* for dental cell isolation studies. The scRNA-seq studies carried out in mouse teeth use a range of enzymes, usually collagenases, that vary according to research group. The Ono group prefers Liberase (Sigma Aldrich) which is a combination of Collagenase I and II and Thermolysin (Nagata et al. 2021; Ono et al. 2016; Takahashi et al. 2019). The Klein group uses Collagenase II (Sharir et al. 2019). The Chai group uses Collagenase I (Jing et al. 2022), the Krivanek group used Collagenase P, or Collagenase D plus Dispase (Krivanek et al. 2020). The Mitsiadis group used Collagenase P for human teeth (Pagella et al. 2021). We found that there were groups of cells that were preferentially included or excluded depending on the enzyme conditions. In particular, the dental mesenchyme cluster was almost entirely isolated using the warm Collagenase D/Dispase II mixture. Almost all T-lymphocytes were excluded by *B. lichenoformis* protease which could be a useful way to reduce the number of white blood cells in the isolation procedures. We used the same labeling system as the mouse studies (10X Genomics) and the same type of sequencer. Nevertheless, the quality of data from the gecko scRNA-seq was lower than the mouse particularly when the warmer temperatures were used. Therefore, the cold method is recommended to supplement the more common protease methods used in mouse or human studies.

We were expecting to find groups of progenitor cells in the scRNA-seq data however due to their low numbers these cells were not encountered. It is possible that some of the dental epithelial and mesenchymal cells contained these putative stem cells. Our previous in situ hybridization experiments on adult geckos located expression of stem cell marker *IGFBP5* primarily in the dental mesenchyme (Handrigan et al. 2010). In agreement, we found *IGFBP5* in a subset of dental mesenchyme cells. The other markers we had studied included WNT pathway transcription factors *TCF3* and *TCF4* both of which were expressed mainly in the dental epithelium. *LGR5* was minimally expressed in the dental epithelium as was *DKK3*. None of these genes were identified in the present data. Further validation of gene expression patterns using in situ hybridization or immunostaining is needed to increase our understanding of the adult gecko stem cells that contribute to new tooth formation.

### Genes and cells required for tooth resorption identified in the gecko bulk RNA-seq data

Currently we know little about how gecko teeth are replaced other than some of the key steps are different than in a mammal. In rodent teeth, the best characterized system, teeth erupt by first creating an eruption pathway through the bone. Osteoclasts under the control of TNFSF11 (RANK ligand or RANKL) and CSF1 are attracted by the dental follicle above the crown of the tooth. The osteoclasts create an eruption pathway allowing the immature tooth to relocate closer to the oral cavity (Nagata et al. 2020; Wise 2009; Wise and King 2008). The pre-eruptive phase of tooth eruption also requires activity of the PTH pathway mediated by the interaction of PTHLH and the PTH1R receptor (Nagata et al. 2020; Philbrick et al. 1998; Richman 2019; Takahashi et al. 2019). The odontoclasts of geckos do express the same markers as osteoclasts (RANKL, CTSK) in humans (Iglesias-Linares and Hartsfield 2017; Wang and McCauley 2011). These molecular profiles of odontoclasts involved in physiological tooth resorption are an extension of the TRAP staining being reported in resorbing human deciduous molars (Sahara et al. 1996; Takada et al. 2004). It is clear that in the gecko there is minimal delay between when a tooth is lost and when the successor appears in the mouth (usually about 3-6 days; (Brink et al. 2022; Grieco and Richman 2018). The clearing of an eruption pathway is replaced by resorption of the functional teeth just at the time when the successional tooth is ready to emerge into the oral cavity. There is similar type of resorption taking place in crocodilians although detailed characterization of the cells involved has not been carried out (Wu et al. 2013). There is one other study on adult lizards and where TRAP+ cells were inside resorbing functional teeth but not in developing teeth (Fuenzalida et al. 1999). No molecular characterization was carried out except for demonstration that selective lectins bind to odontoclasts. Here we found the cellular basis for this quick eruption is the timed invasion of odontoclasts. This is supported by the profiling of transcripts in the functional teeth that confirm expression of the typical genes involved in the induction and activation of osteoclasts. There is conservation of the resorptive process paving the way for functional studies in the future.

An unexpected molecular finding was the discovery of chemotaxis genes in the functional teeth and developing teeth. The expression of the *SEMA3A* gene is of particular interest and may play a dual role in recruitment of odontoclasts and axon guidance. Thus far the data in mouse studies suggests that *SEMA3A* is expressed in the osteoclasts that are creating the eruption pathway (Yu et al. 2019). In addition, the SEMA3A receptors, Neuropilin 1 and Plexin A1 are also expressed during tooth eruption (Sijaona et al. 2012). Further in vivo experiments are needed to test the role of SEMA3A, CTSK and RANKL in directing tooth resorption in the gecko.

### Molecules that are associated with the dental follicle and tooth eruption are expressed in gecko

Tooth eruption is a functional requirement of the dentition across the entire animal kingdom. In squamate reptiles (snakes and lizards), the newly forming teeth initiate deep in the jaws, in a tooth forming field where tissue interactions are taking place. After tooth initiation, the squamate reptiles have teeth that erupt towards the oral cavity while remaining attached to the dental lamina (Brink et al. 2021; Handrigan et al. 2010). This mechanism is different in crocodilians where the dental lamina is discontinuous and found within a tooth socket containing the functional tooth (Wu et al. 2013). Humans (Ooë 1981) and other diphyodont mammals (Buchtova et al. 2012) also have an early separation of the successional tooth from the dental lamina which degrades via apoptosis.

Another difference between mammals and lizards is that the mammal absolutely requires the dental follicle for tooth eruption (Cahill and Marks 1980; Larson et al. 1994; Marks and Cahill 1984; 1987). The dental follicle is a sac of mesenchyme cells that surrounds the tooth, separating it from the bone. The follicle later creates the tooth bone interface which is mammals consists of the periodontal ligament and cementum (Nagata et al. 2022). Upon comparing the mouse data dental follicle transcriptome data (Jing et al. 2022; Takahashi et al. 2019) to the gecko we were interested to find cell types that express genes characteristic of the dental follicle. The dental mesenchyme (cluster 3) expressed *SFRP2, IGFBP5, FOS* and *SGK1*, all of which are restricted to the mouse dental follicle (Hermans et al. 2022). It is possible that these *SFRP2*+ cells are dental follicle progenitors. Interactions mediated via PTHRP signaling are needed to maintain the follicle, to create a periodontal ligament attachment and to permit tooth eruption (Takahashi et al. 2019). *Pthlh* was expressed in the mouse dental follicle (Takahashi et al. 2019) but in the gecko there was no expression of *PTHLH* in the putative dental follicle (as marked by *SFRP2* expression). Since geckos and all other dentate reptiles except crocodilians, lack a periodontal ligament (Bertin et al. 2018; LeBlanc et al. 2021), *PTHLH* may take on different functions in geckos, the most likely of which is to recruit odontoclast precursors. *PTHLH* is expressed in the stellate reticulum of the second generation tooth which is next to the TRAP+. CTSL+ positive odontoclasts.

There is at least one more member of the SFRP family, *SFRP1*, present in developing teeth so there could be some functional redundancy. There are significant phenotypes in the head and body axis of the *Sfrp1/Sfrp2* homozygous null animals but there are no reports on the dental or tooth eruption phenotypes of the compound knockouts (Satoh et al. 2006).

In conclusion the comparison of transcriptomes in a polyphyodont reptile to those of the mouse have already revealed stark differences. Our studies suggest that there has been independent (convergent) evolution of the trait of tooth eruption in lizards and snakes. Moreover, the second generation teeth likely do communicate with the functional teeth as shown by our work. We removed second generation teeth and followed the eruption process for many months. The functional teeth were retained longer in sites where there were no immediate successors (Brink et al. 2022). Thus instead of an eruption pathway as in the mammal, the gecko and other squamate reptiles use the enamel organ of the successional teeth to trigger tooth resorption of the functional teeth, thus creating an eruption pathway. New molecules such as SEMA3A may also participate in this process. The profile of stem cells differs from the mouse incisor. Future studies on the gecko will uncover the molecular basis of convergent evolution in the dentition.

## METHODS

### Total RNA extraction for bulk RNA sequencing and qRT-PCR

We removed functional and developing, unerupted teeth and these were pooled into three biological replicates for bulk RNA sequencing (Fig. S1A). There were 3 functional or 3 developing teeth pooled for each biological replicate. For the qRT-PCR, independent biological replicates were collected and each contained 10-12 teeth. All teeth were snap frozen in liquid nitrogen. Each sample was disrupted using beads (Buffer RLT from Qiagen microRNA kit, Omni bead Ruptor 96, Lysing matrix E) followed by a Qiagen Shredder column (ID:79656). Total RNA was extracted using a Qiagen RNeasy micro kit (ID: 74004). RNA quality was determined with Agilent Bioanalyzer 2100. Samples with an RNA Integrity (RIN) greater than 7 were sequenced using Illumina Nextseq 2000 or used for qRT-PCR (Table 1).

**Table 1.**
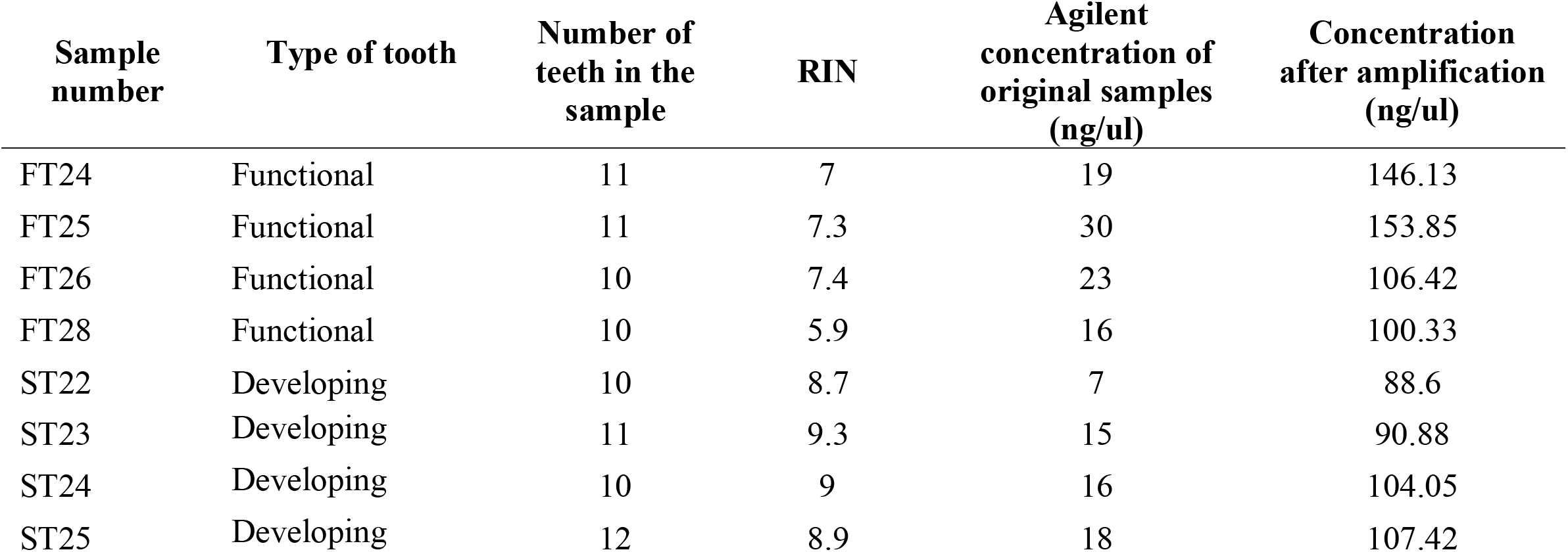
Characteristics of biological replicates used for qRT-PCR

| **Sample number** | **Type of tooth** | **Number of teeth in the sample** | **RIN** | **Agilent concentration of original samples (ng/ul)** | **Concentration after amplification (ng/ul)** |
| --- | --- | --- | --- | --- | --- |
| FT24 | Functional | 11 | 7 | 19 | 146.13 |
| FT25 | Functional | 11 | 7.3 | 30 | 153.85 |
| FT26 | Functional | 10 | 7.4 | 23 | 106.42 |
| FT28 | Functional | 10 | 5.9 | 16 | 100.33 |
| ST22 | Developing | 10 | 8.7 | 7 | 88.6 |
| ST23 | Developing | 11 | 9.3 | 15 | 90.88 |
| ST24 | Developing | 10 | 9 | 16 | 104.05 |
| ST25 | Developing | 12 | 8.9 | 18 | 107.42 |

### Bulk RNA sequencing and data analysis

Qualifying samples were prepped following the standard protocol for the NEBnext Ultra ii Stranded mRNA (New England Biolabs) at Biomedical research Center sequencing Core at University of British Columbia. Sequencing was performed on the Illumina NextSeq2000 with Paired End 59bp × 59bp reads. Sequencing data was demultiplexed using Illumina’s bcl2fastq2 and approximately 20 million read pairs were obtained for each sample. The bulk RNA-seq data was deposited into the National Center for Biotechnology Information Gene Expression Omnibus database (Accession GSE220653).

As previously described in Vuilleumier *et al* (Vuilleumier et al. 2019), multiple differential expression analysis pipelines were applied and the results combined. In the current study the pipelines primarily used the draft reference genome and transcriptome from Xiong *et al* and http://gigadb.org/dataset/100246. The important gene *CTSK* was however missing from the published annotations so we therefore called genes *de novo* with the code BRAKER ; (Barnett et al. 2011; Bruna et al. 2021; Buchfink et al. 2015; Hoff et al. 2016; Hoff et al. 2019; Li et al. 2009; Lomsadze et al. 2014; Stanke et al. 2008; Stanke et al. 2006) and https://github.com/Gaius-Augustus/BRAKER) using our RNA-seq read alignments as hints. We then extracted the *CTSK* gene structure and associated transcripts from those results and added the information to the reference genome/transcriptome before doing the differential expression analysis (Table S2). Only genes achieving a false discovery rate of 0.05 in at least half of the ten RNA-seq pipelines applied were considered as significantly differentially expressed. Volcano plots for some of the individual analysis pipelines were created using the R package Enhance Volcano plot (https://github.com/kevinblighe/EnhancedVolcano).

Several genes annotated as “unknown” in the published genome were searched using blastx against the nucleotide collection (nr/nt) (Camacho et al. 2009; Sayers et al. 2022) in order to assign more meaningful gene names. The gene names were changed to their respective human orthologue to improve the quality of post-hoc analyses. To determine gene ontology of the main biological processes in the two types of teeth, the list of differentially expressed genes was entered into the Panther Classification system (Pantherdb.org; Table S4).

### qRT-PCR validation of bulk RNA-seq

Genes were selected for validation with qRT-PCR with a log_2_ fold change of at least 0.9, a false discovery rate lower than 0.05 and/or biological relevance for tooth development and osteoclast differentiation and function. Ten nanograms of total RNA from each replicate was amplified for 2h with the Qiagen QuantiTect whole transcriptome amplification kit (Table 1). Aliquots were taken before inactivating Repli-g Midi DNA polymerase and used to quantify double stranded cDNA with Quant-iT™ PicoGreen™ kit (Thermo-Fisher). Sybr green-based quantitative qRT-PCR was conducted using a BioRad kit (#1725271; SsoAdvanced Universal SYBR-Green supermix). Primers were designed based on the gecko genome and transcriptome sequence (Table S5). Quantitative-PCR cycling conditions were: 95°C for 10min, 40X (95°C for 5s, 60°C for 20s). Gene expression levels were calculated using ∆CT method with normalization against 18s ribosomal RNA (Applied Biosystems #4318839). All qRT-PCR reactions were performed in triplicate for each sample, four biological replicates were analyzed for each type of tooth (Table 1). The average of 2^-^∆^CT^ of functional teeth samples was computed and used to normalized the data of each sample. T-test were computed between the normalized 2^-^Δ^CT^ of functional and developing teeth samples to determine statistically significant differences in expression between.

### Single Cell RNAseq

In order to identify cell subpopulations in the odontogenic regions of the jaws, we dissected tissues enclosing oral mucosa, dental lamina, dental mesenchyme, immature cap and bud stage teeth and bell stage, mineralizing teeth (Fig. S1B). Tissues were dissected and placed in cold 1X HANKS solution without Calcium and Magnesium and 10% fetal bovine serum (FBS, Gibco). The tissues were chopped into small segments. Then, they were placed in the dissection media with collagenase D/ Dispase II (SIGMA cat no. 11231857001/SIGMA cat no. D4693) at 37°C for 4-5 hours in an orbital shaker. Alternatively, the tissue from the second animal was dissociated with a cold temperature enzyme protease from *Bacillus lichenoformis* (cat no. Sigma p5380) for 6 hours at 4°C. Once the cells were dissociated, cells were spun down and counted. They were transferred into FACS buffer (2 mM EDTA, 2% FBS, 1X PBS) and resuspended in 2 ml final volume. Cells were passed through a 70 µm cell filter to remove clumps. Live versus dead cells were distinguished with propidium iodide exclusion (1:5000 in FACS buffer). Cells were sorted in DMEM + 5% FBS media in a Beckman Coulter Cytoflex LX Analyzer. Cells were concentrated to 50,000 cell/ml. The cell suspension was loaded on the 10x genomics single cell controller for capture in droplet emulsion. Libraries were prepped using the Chromium Single Cell 3’ Reagent v3 Chemistry kit and the standard protocol was followed for all steps. Libraries were then sequenced on an Illumina Nextseq500. 10x genomics Cell Ranger 2.0 was used to perform de-multiplexing, alignment, counting, clustering, and differential expression. The scRNA-seq raw and processed data were deposited to the National Center for Biotechnology Information Gene Expression Omnibus database (GEO; https://www.ncbi.nlm.nih.gov/geo/; accession number GSE221110). Cellranger was used to merge cells data coming from the same animals. Cellranger exports were further analyzed using - Seurat package 4.0.4. A differentially expressed gene list for each cluster was computed using the Findmarkers function (Table S2). Note that the mitochondrial genes in the gecko are not annotated so we did not filter on high levels of mitochondrial DNA in the quality control steps.

### Immunofluorescence and TRAP staining

In order to identify odontoclasts, tartrate acid resistant phosphatase (TRAP) staining was carried out on paraffin sections of gecko tooth families. Slides were placed in pre-warmed TRAP solution (0.92 % Sodium Acetate Anhydrous, 1.14%, Tartaric Acid, 0.01% Napthol AS-MX Phosphate and 0.06% Fast Red Violet LB Salt; Sigma Aldrich) and incubated at 37°C for 60 mins. Slides were rinsed in distilled water, counterstained with 0.02% Fast green for 1 minute and then coverslipped.

Immunofluorescence staining was carried out on paraformaldehyde fixed tissues embedded in paraffin (Brink et al. 2021). The primaries antibodies were: anti-Cx43 (Sigma-Aldrich, C6219, 1:1000); anti-RANKL (Abcam, ab65024, 1:200); anti-CTSK (Abcam, ab37259, 1:200); anti PITX2 (R&D systems, AF7388, 1:500). Antibodies to Cx43, RANKL and CTSK were diluted in 10% goat serum, 1XPBS, and 0.1% Tween-20. PITX2 was diluted in 5% bovine serum, 0.1% Tween-20 in Tris buffered salt solution. The secondary antibodies were diluted to 1:250 (Goat anti-rabbit IgG Cy5; Invitrogen A10523; goat anti-mouse IgG AlexaFluor 488, Invitrogen A11029; goat anti-rabbit IgG AlexaFluor 488, Invitrogen A11034). Nuclei were stained with Hoechst 33258 (Sigma, 10 mg/ml; 30 min). Slides were scanned in the fluorescence mode using a 3DHISTECH Slide scanner and a 20X objective (3DHistech.com, Hungary).

### RNAscope in situ hybridization

We selected genes for validation with RNAscope from the bulk RNAseq data based primarily on biological relevance, low FDR and log_2_ fold change of at least 0.9 (functional or developing tooth samples, *SEMA3A, SEMA3E, PTHLH*). Flanking 5’ and 3’ UTRs plus coding sequences for each of the genes were extracted from the gecko genome in our lab. In order to find the UTRs, we used BED tools to extract the coding sequences and UTRs (Quinlan and Hall 2010). We validated the sequences with the IGV browser downloaded from the Broad Institute (Robinson et al. 2011). For each selected gene, we next confirmed that there was little homology to other related family members using nr Blast against *Gekko japonicas* (Camacho et al. 2009). In the case of *COL1A1*, areas of very highly conserved sequence between *COL1A1* and *COL1A2* were avoided. These trimmed sequences were provided to the company ACDBio who designed custom DNA probes. The probes used are: Emac-SEMA3A-C1 (ref 1099931-C1), Emac-*SEMA3E*-C1 (ref 1099911), Emac-*PTHLH*-C1 (ref1208601-C1), *COL1A1*, Emac-*COL1A1-C1* (ref 1214171-C1). The following ZZ pairs were used: Emac-SEMA3A,18ZZ probe targeting 47-912 of the provided sequence; 20ZZ probe named Emac-SEMA3E targeting 344-1312 of the provided sequence; 10ZZ probe named Emac-PTHLH targeting 16-525 of the provided sequence; 20ZZ probe named Emac-COL1A1 targeting 251-1133 of the provided sequence and Emac-SFRP2, 20 ZZ probe targeting 344-1496 of the provided sequence. *COL1A1* was used as a positive control for the in situ hybridization conditions. The RNAscope 2.5 HD detection red kit was used according to manufacturer indications with the following modifications: amplification step 5 was carried out for 50 minutes for all probes except for *PTHLH* which was carried for 90 minutes. Slides were coverslipped and scanned with a 3DHISTECH Slide scanner (3DHistech.com, Hungary) using a 20X objective with brightfield illumination.

## Supporting information

Henriquez Table S1

Henriquez Table S2

Henriquez Table S3

Henriquez Table S4

Henriquez Table S5

## Acknowledgements

The authors would like to thank the Biomedical Research Sequencing core at UBC (Tara Stach and Ryan Vander Werff), Preety Panwar and Dieter Bromme for providing initial reagents to stain odontoclasts and Mark Sgro for help with blasting the gecko sequences.

