## Supplementary figures for "Transcriptomic profiling of the adult reptilian dentition sheds light on the genes regulating indefinite tooth replacement"

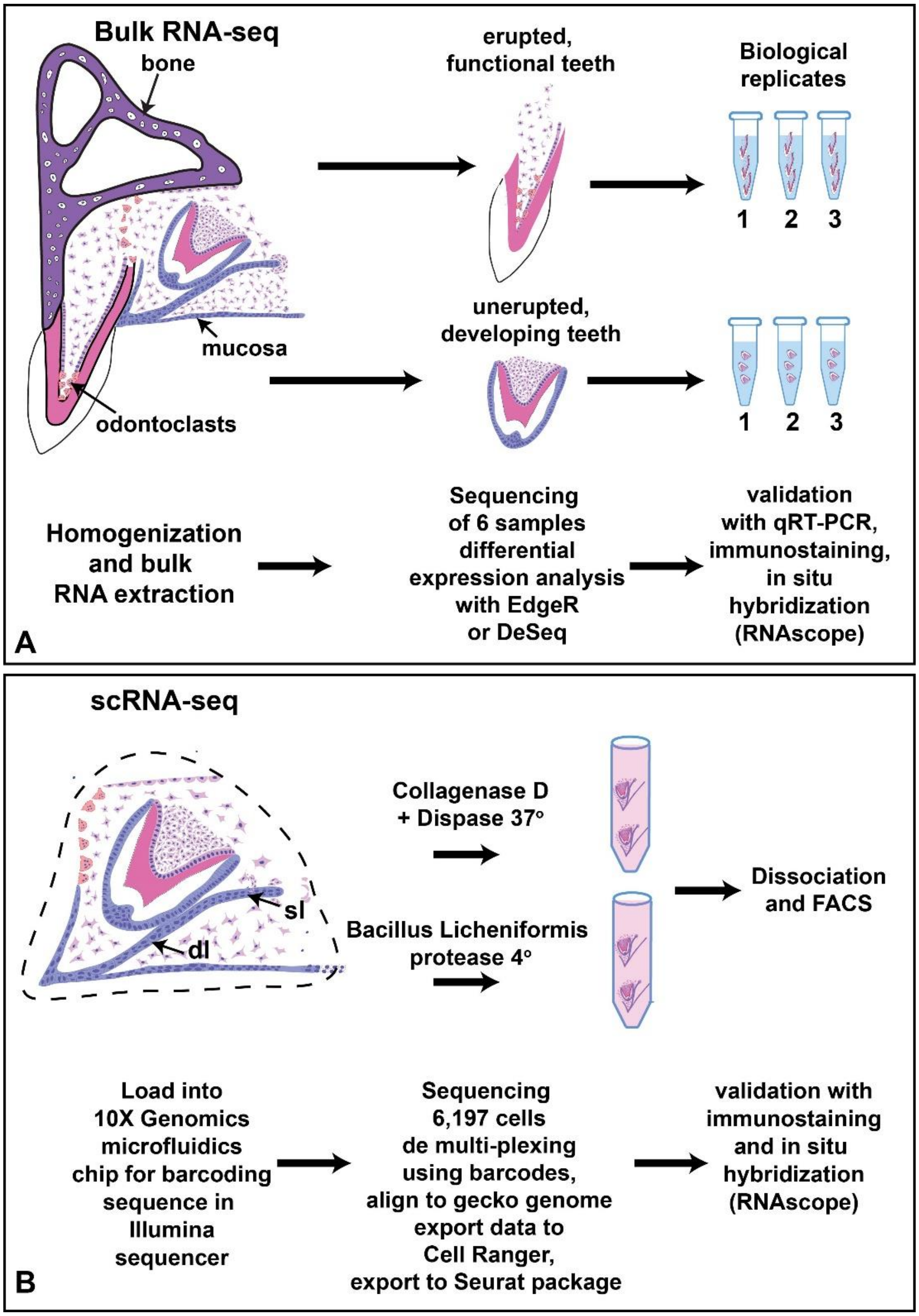


### Figure S 1

Diagram depicting the pipeline used to characterize the odontogenic tissues of the adult leopard gecko. Functional teeth with signs as well as developing teeth, were surgically extracted from adult animals. Total RNA was extracted from each type of samples, sequenced, and a differentially expression between functional and developing teeth was computed. Candidate transcript relevant for the process of tooth development, for clastic cells activity or recruitment and previously uncharacterized genes were validated through qRT-PCR and RNAscope (A). Oral mucosa encompassing odontogenic tissues, oral epithelium, and other associated structures were extracted from two animals. Cells were dissociated with two different enzymatic treatments. Individual alive cells were FACS sorted and submitted to single cell RNA seq to identify cell populations associated with postnatal continuous tooth replacement in these animals, cells belonging to developing teeth, cells responsible for continuous tooth shedding (B).

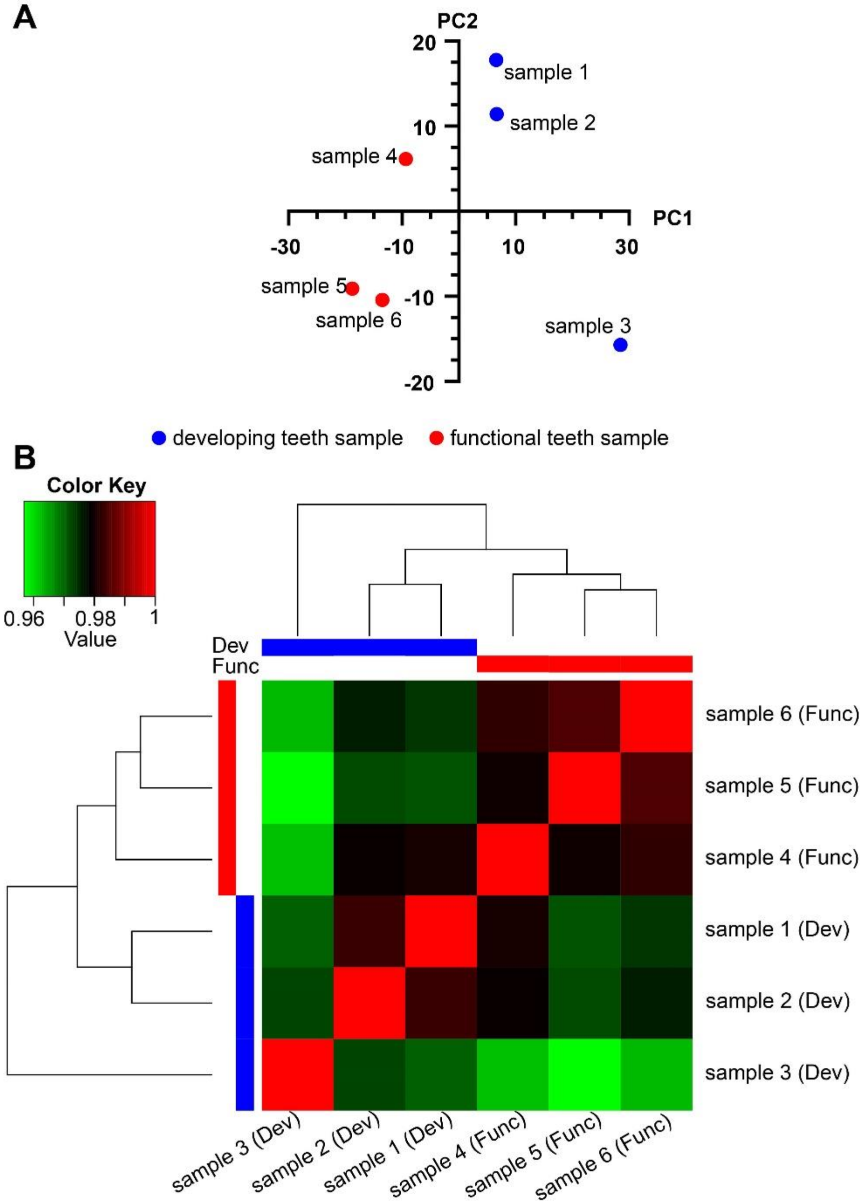


C

| sample | animal | type |
|---|---|---|
| sample 1 | gecko 1 | developing |
| sample 2 | gecko 1 | developing |
| sample 3 | gecko 2 | developing |
| sample 4 | gecko 1 | functional |
| sample 5 | gecko 2 | functional |
| sample 6 | gecko 2 | functional |

### Figure S 2

Quality control figure of bulk RNAseq showing how samples group across PC1 according of type of samples (functional or developing) (A). Samples group according to animal origin in PC2. (B). Hierarchical clustering of samples according to similarities of the sequenced RNA. (C) Samples identified by animal number.

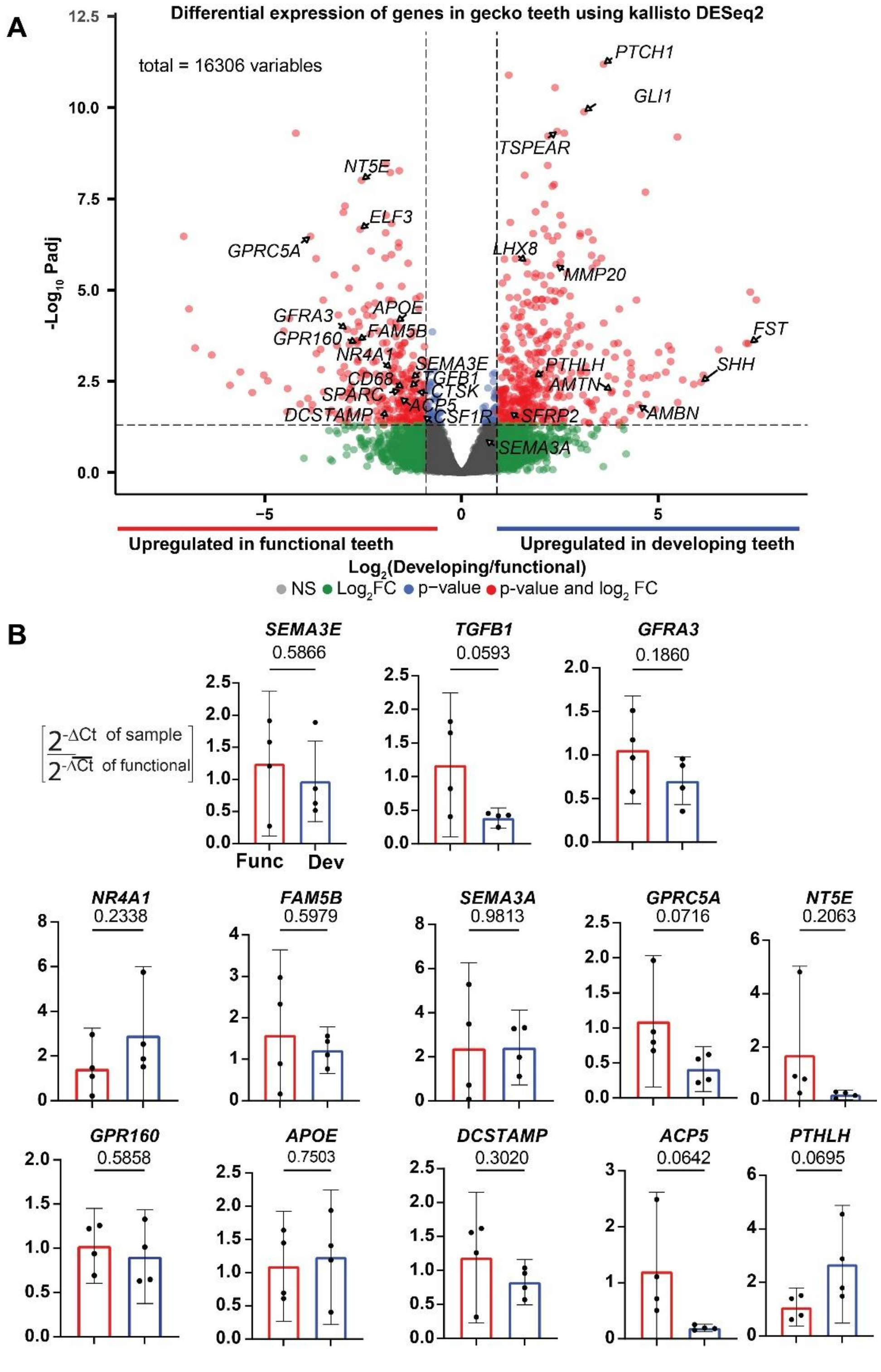


**Figure S 3**

Volcano plot of transcripts sequenced in bulk RNAseq. Adjusted p-value and enrichment levels for this specific plot where computed with Deseq2. Dots in red are transcripts with a p-value over 0.05 and a $Log_2$ of the ratio between the expression level of developing and functional teeth over ± 0.9 (A). B) $2^{-\Delta CT}$ normalized by the average $2^{-\Delta CT}$ of functional teeth of transcripts that were determined to be statistically significantly expressed between functional and developing teeth samples in bulk RNAseq, in any enrichment analysis, but failed to present a differentially expressed ΔCT between functional and developing teeth when tested in independent samples using qRT-PCR. 95% confident interval are plotted (B).

Figure S3

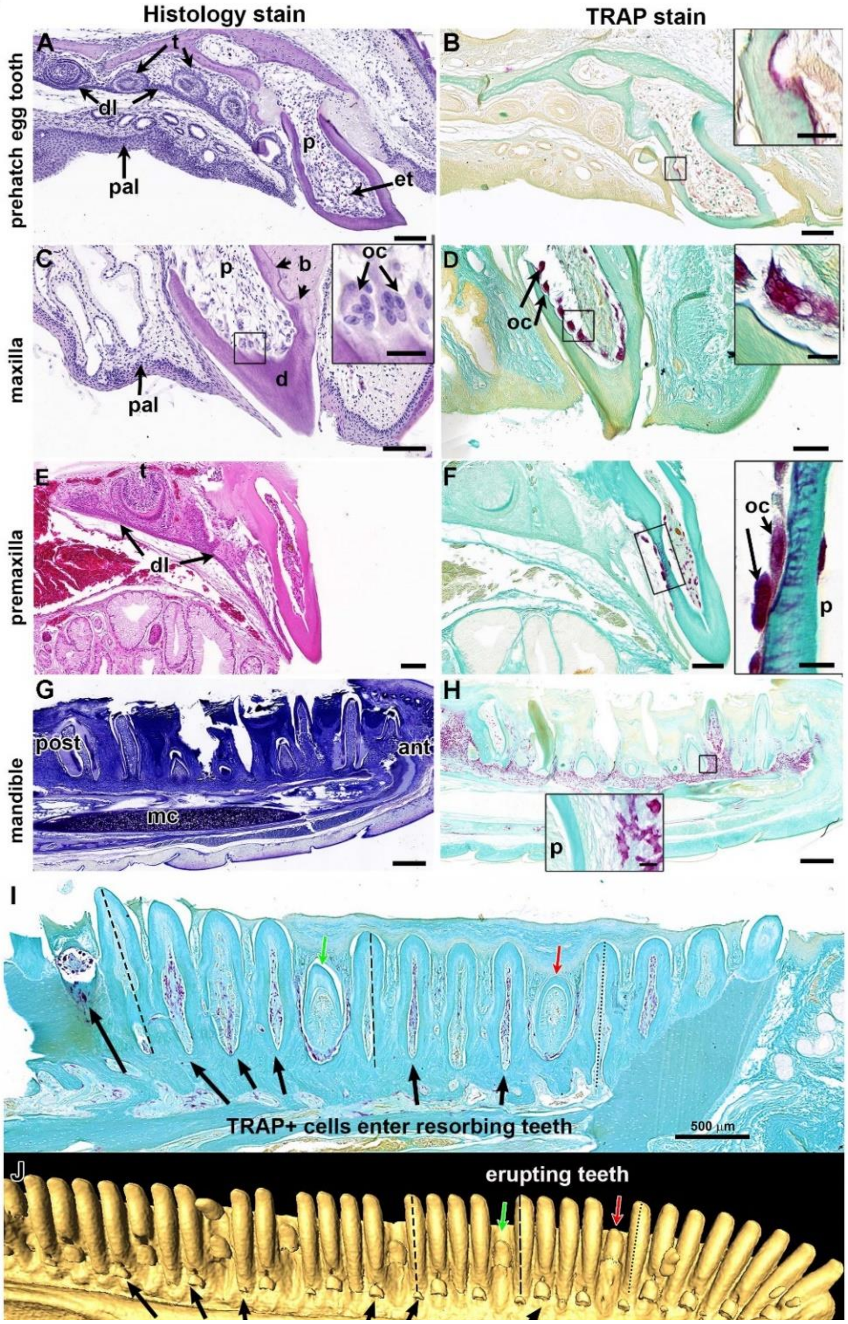


### Figure S 4

Tooth succession correlates with invasion of odontoclasts (A) A prehatching animal showing the egg tooth and 5 replacement teeth connected by an epithelial dental lamina. (B) Near-adjacent section showing TRAP+ cells in the egg tooth (B, inset). (C) Transverse section of an adult maxilary tooth. Dentin is directly attached to the bone (arrowheads). Multinucleated odontoclasts have replaced odontoblasts (inset). (D) Near-adjacent section showing multinucleated cells are TRAP+ . (E) Transverse section of a premaxillary teeth. (F) Near-adjacent section showing TRAP + cells attached to the outer and inner surface of the dentin. Signal is inside the dentinal tubules (inset). (G) Sagittal section of a mandible showing teeth in different stages of development. Notice that Meckel's cartilage persists in adult geckos. (H) Near-adjacent section with TRAP+ positive cells inferior to the teeth and in some cases extending into the pulp chamber. There is no TRAP staining in pulps of developing teeth (inset). I) Trap+ cells entering some functional teeth (black arrows). Developing teeth have no TRAP+ cells (Red and green arrows). J) µCT of same jaw as in I showing the same erupting teeth (green and red arrows). Unerupted teeth are developing apical to the functional teeth (black arrows). Key: ant – anterior, b – bone, d – dentin, dl – dental lamina, et – egg tooth, mc – Meckel's cartilage, oc – odontoclast, p – pulp, pal – palate, post-posterior.

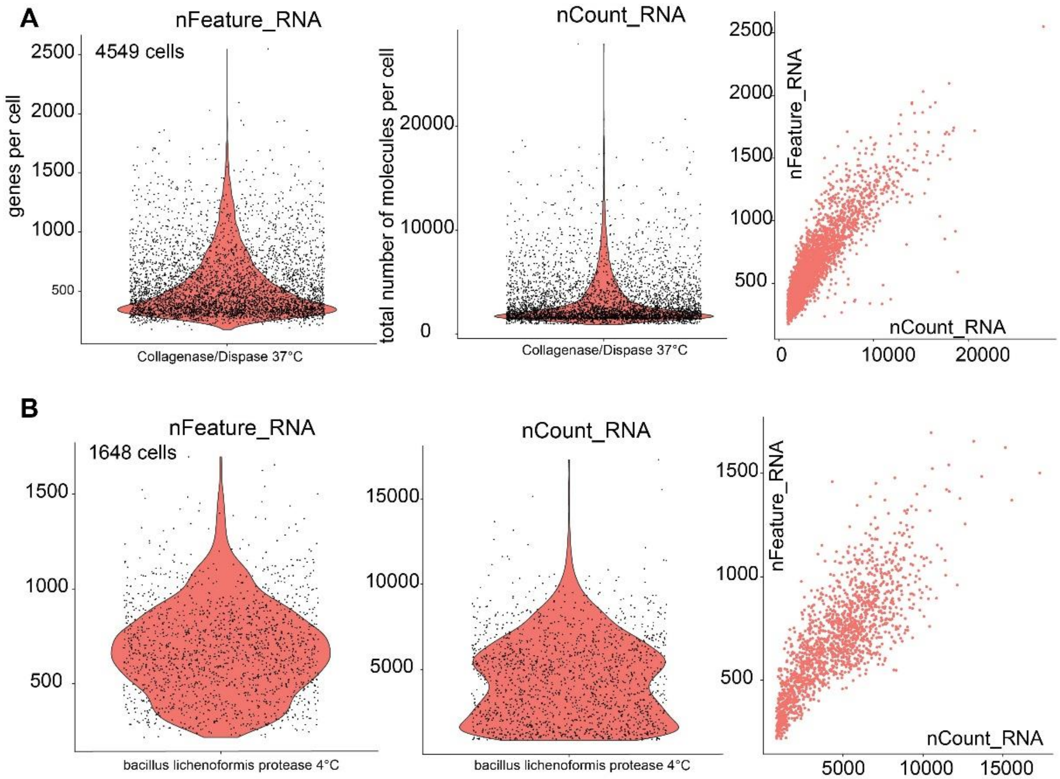


C

| | collagenase/Dispase - 37°C | Bacillus *Lichenoformis* - 4°C |
|---|---|---|
| Estimated Number of Cells | 4,549 | 1,648 |
| Fraction Reads in Cells | 75.70% | 90.70% |
| Pre-Normalization Mean Reads per Cell | 99,510 | 237,554 |
| Post-Normalization Mean Reads per Cell | 73,107 | 190,373 |
| Median Genes per Cell | 461 | 665 |
| Median UMI Counts per Cell | 2,188 | 4,318 |

### Figure S 5

Features or transcripts read versus genes or counts. A) The cells isolated with collagenase and dispase at 37 °C are compared to B) cells isolated with B. lichenoformis at 4°C. The warm temperature resulted in more transcripts but these were mapped to fewer genes per cell compared to cold-temperature isolation. In contrast the cold-temperature isolation results in transcripts distributed across many more genes. Quality was improved for each cell but fewer cells overall were isolated with the cold enzymatic separation method.

**A**

**Effect of enyzme treatment on contribution of cells to each cluster**

| cluster | cell type | collagenase/ dispase 37°C | *Bacillus lichenoformis* Protease 4°C | total |
|---|---|---|---|---|
| 0 | epithelial | 938 | 673 | 1611 |
| 1 | T-lympocytes | 1279 | 14 | 1293 |
| 2 | unknown | 750 | 100 | 850 |
| 3 | mesenchyme | 569 | 86 | 655 |
| 4 | unknown | 197 | 315 | 512 |
| 5 | cycling cells | 309 | 119 | 428 |
| 6 | dental epithelial | 130 | 198 | 328 |
| 7 | white blood cells | 168 | 27 | 195 |
| 8 | unknown | 131 | 13 | 144 |
| 9 | blood vessel | 53 | 21 | 74 |
| 10 | unknown | 20 | 39 | 59 |
| 11 | unknown | 5 | 43 | 48 |
| | total | 4549 | 1648 | 6197 |

**B**

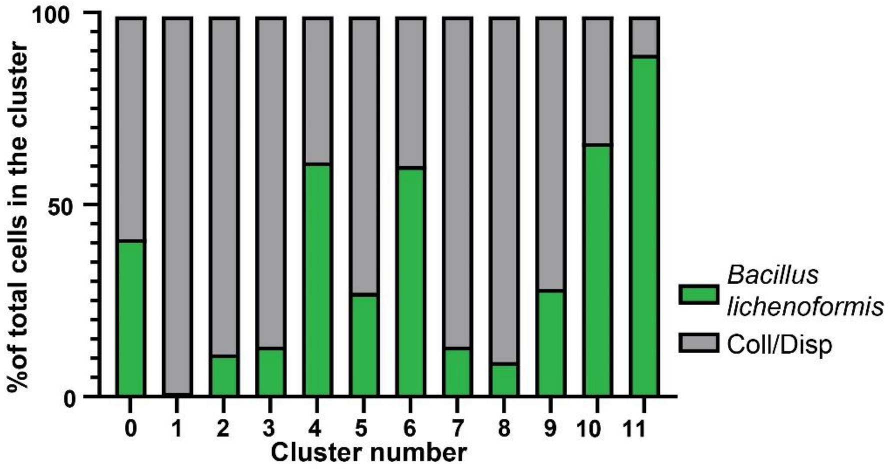


**Figure S 6**

Contribution from enzymatic dissociation methods to each cluster. Total number of cells present at each cluster and number of cells per cluster according to the enzymatic method used to extract them.

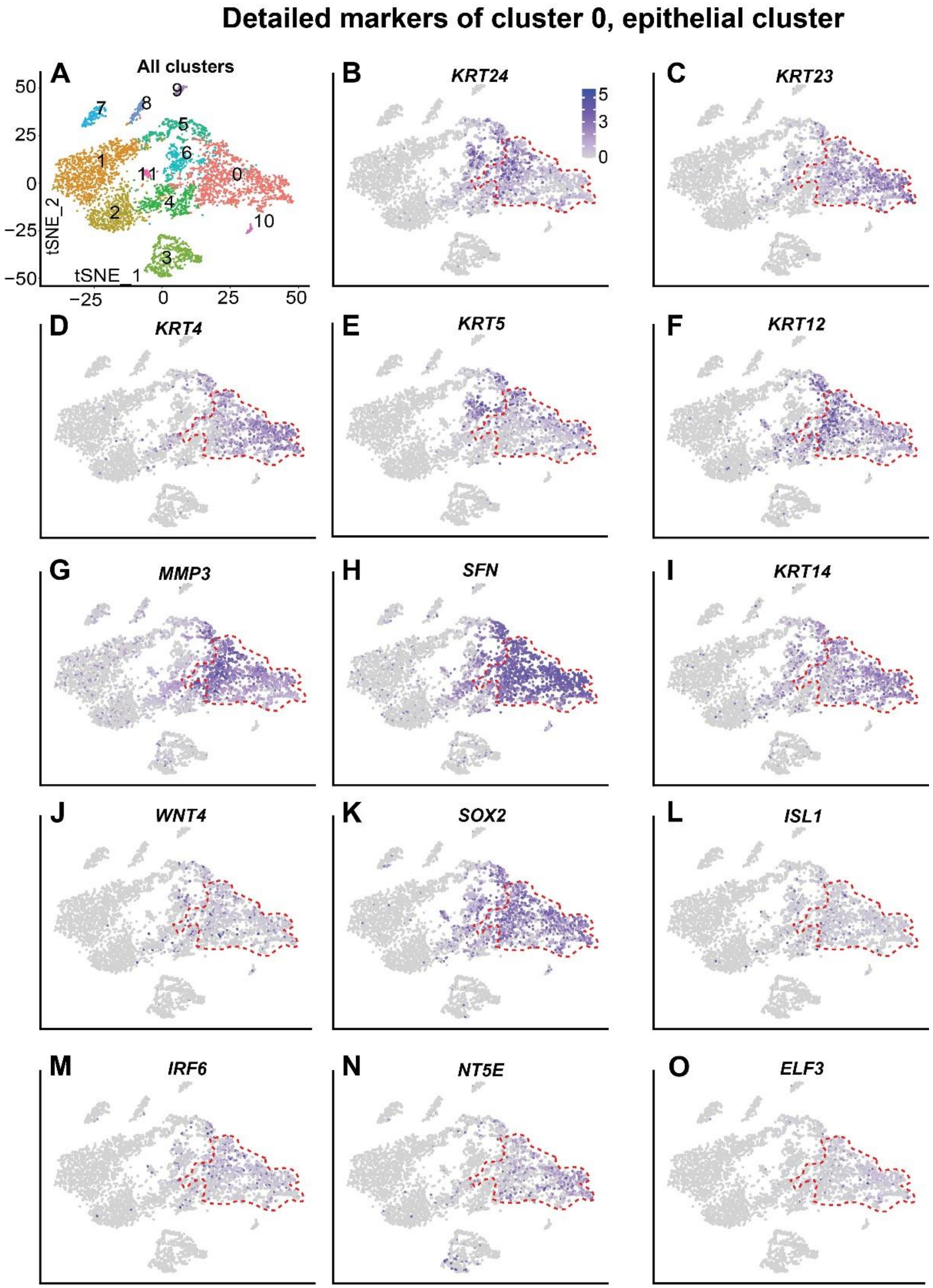


**Figure S 7**

tSNE plots of all the clusters and transcripts differentially expressed in cluster 0. Cluster zero is surrounded by segmented red lines in each tSNE plot for each gene. Transcript name and gene id are indicated in the superior part of each plot.

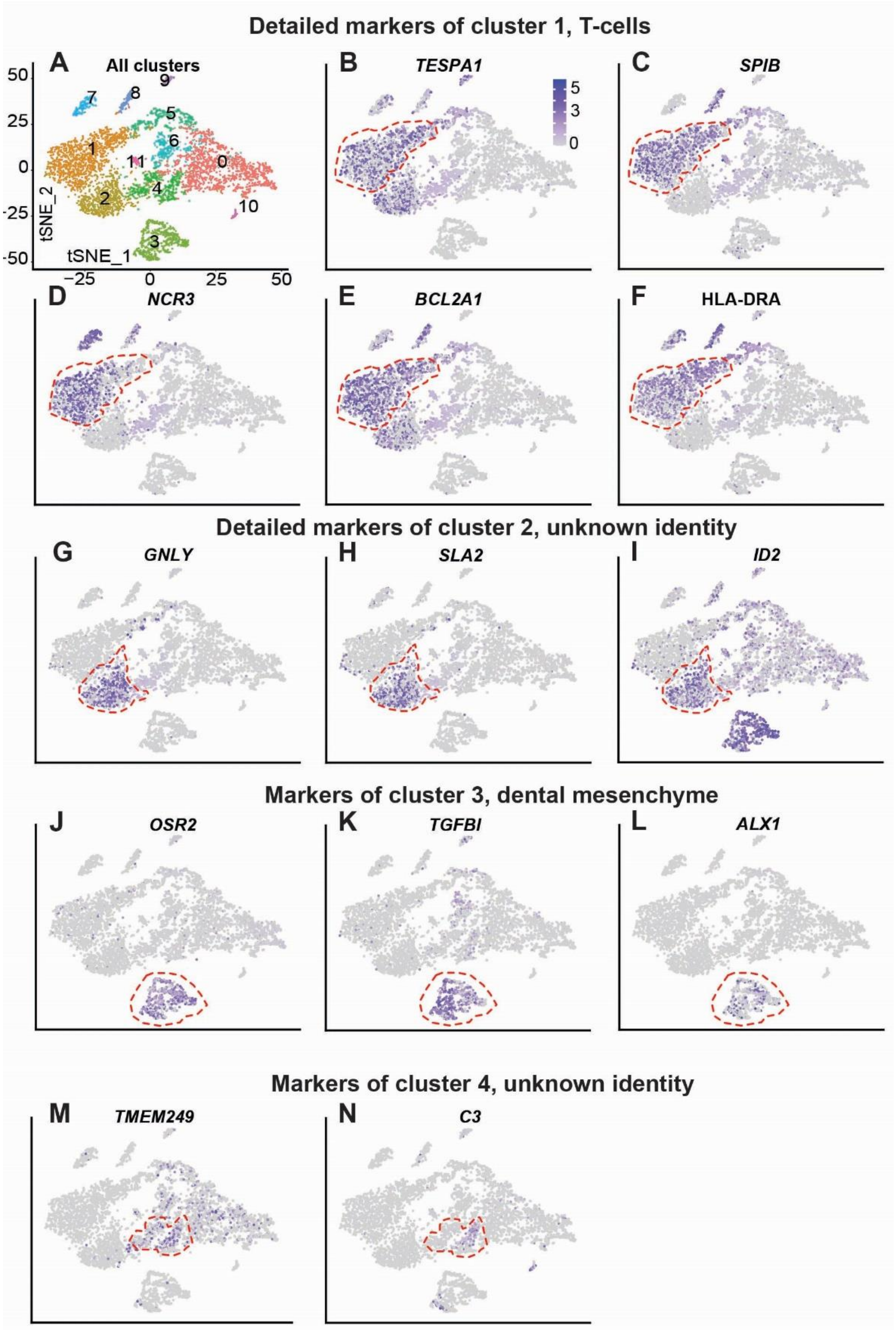


**Figure S 8**

tSNE plots of all the clusters and transcripts differentially expressed in cluster 1, 2, 3 and 4. Respective clusters are surrounded by segmented red lines in each tSNE plot for each genes. Transcript name and gene id are indicated in the superior part of each plot.

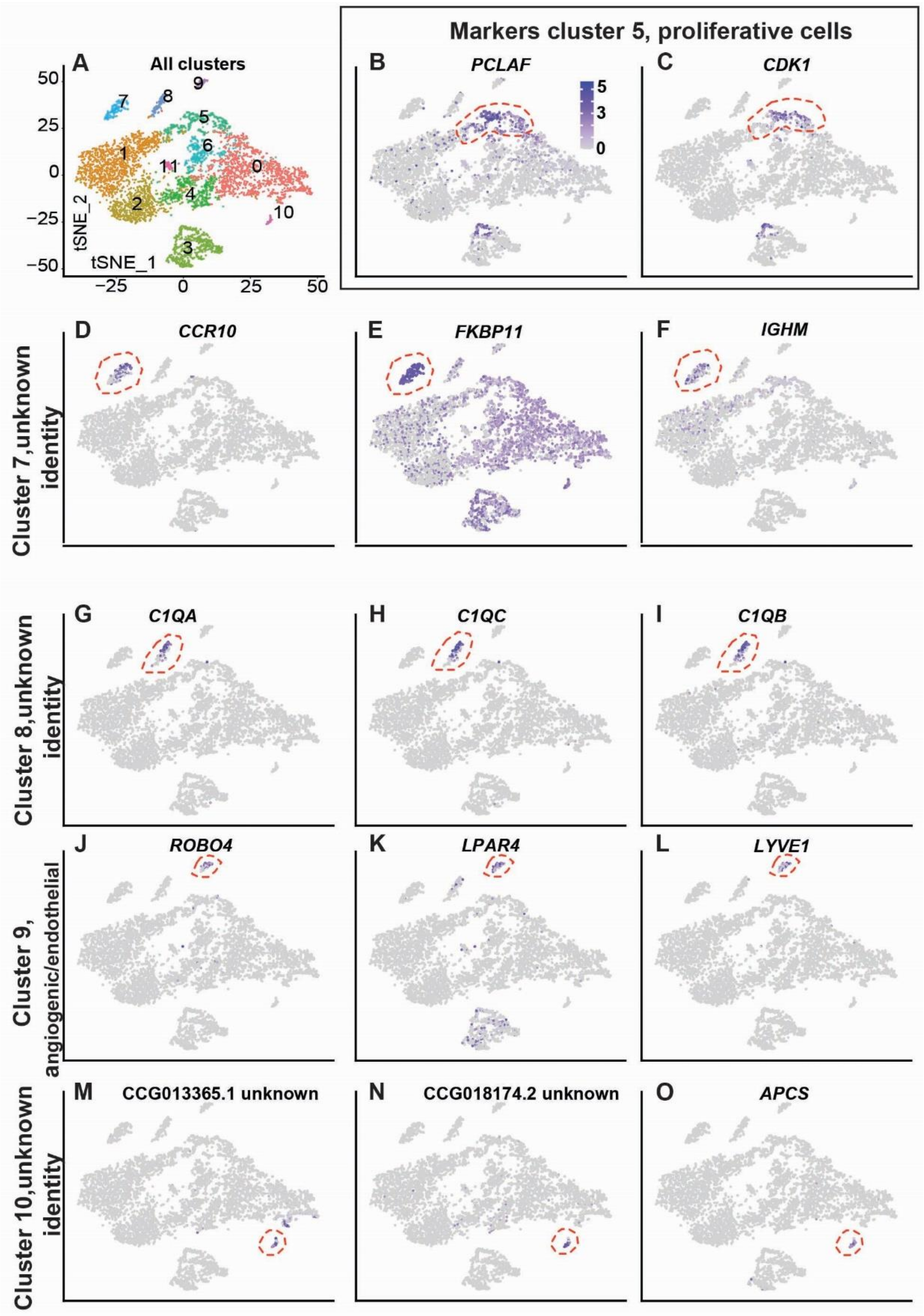


**Figure S 9**

tSNE plots of all the clusters and transcripts differentially expressed in cluster 5, 7, 8, 9 and 10. Respective clusters are surrounded by segmented red lines in each tSNE plot for each gene. Transcript name and gene id are indicated in the superior part of each plot.

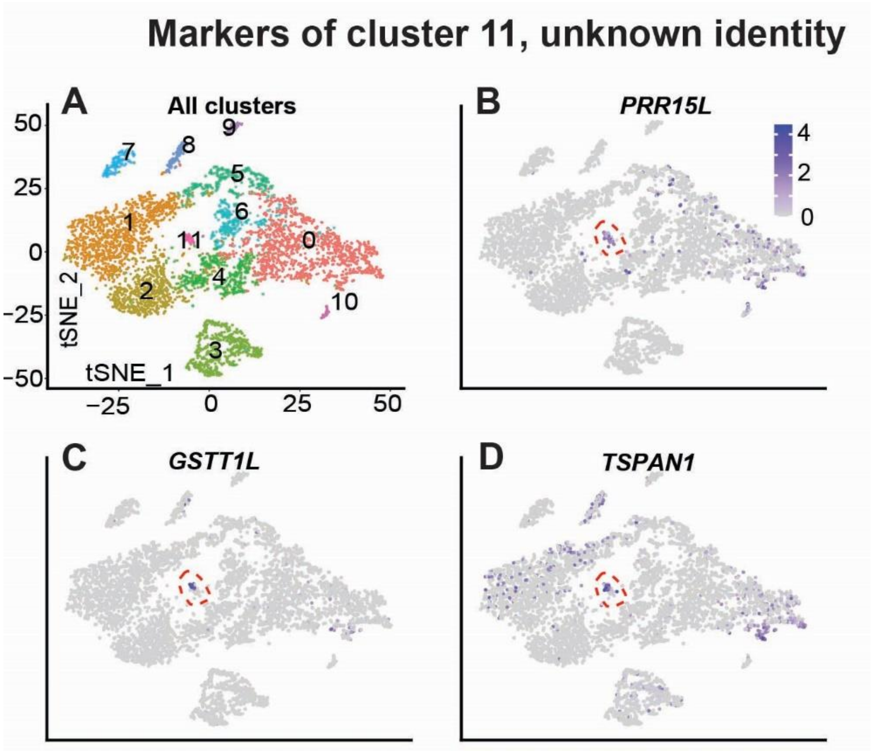


**Figure S 10**

tSNE plots of all the clusters and transcripts differentially expressed in cluster 11. Cluster 11 is surrounded by segmented red lines in each tSNE plot for each gene. Transcript name and gene id are indicated in the superior part of each plot.

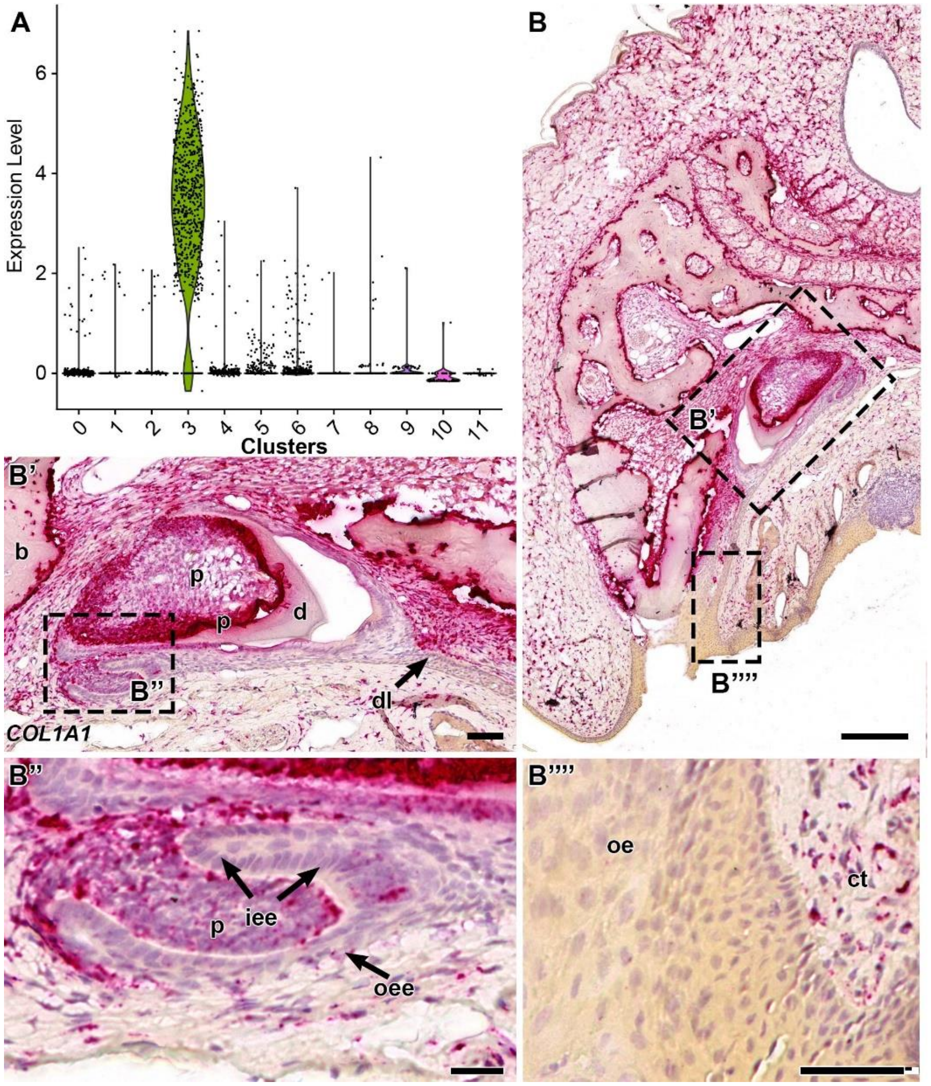


### Figure S 11

Violin plot and spatial location of *COL1A1* transcripts in gecko teeth. A) *COL1A1* was statistically significantly enriched in cluster 3 cells. B-D) RNAscope chromogenic staining of *COL1A1* in maxillary section. C) Strong expression in the mesenchyme of developing teeth but no expression in the epithelial dental lamina. C') Expression of *COL1A1* is limited to the mesenchyme and not the enamel organ. D) The oral epithelium is negative for COL1A1 and the adjacent connective tissue has abundant signal confirming probe specificity. Key: b-bone, ct-connective tissue, d-dentin, iee-inner enamel epithelium, oee- outer enamel epithelium, p- papilla. Scale bar: B= 200 μm; C-D'= 50 μm.

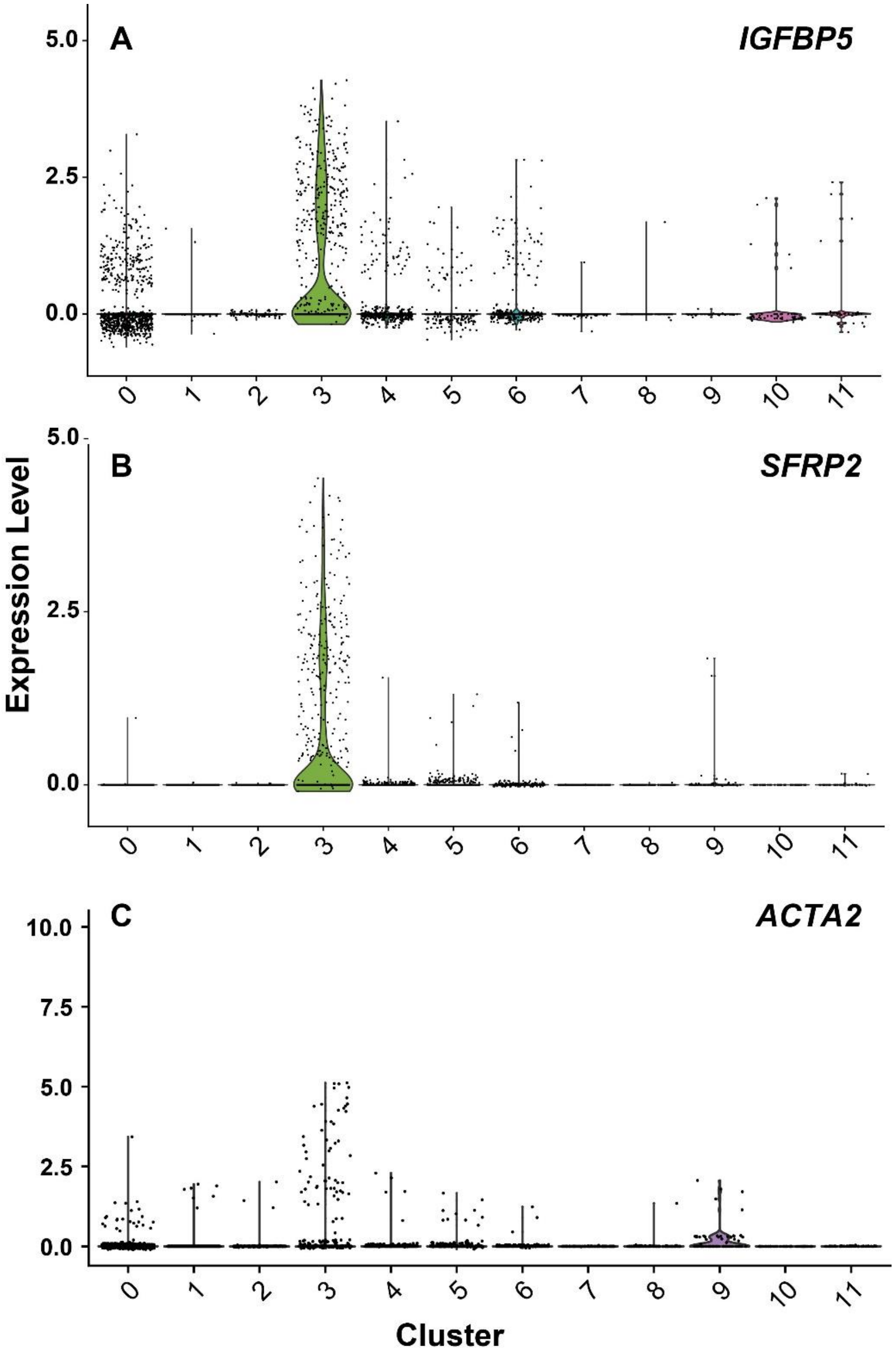


**Figure S 12**

Violin plots for markers of dental follicle. Expression level corresponding to normalized expression. Cluster identity is indicated with numbers in the horizontal axis.
